# Interferons drive development of novel interleukin-15-responsive macrophages

**DOI:** 10.1101/663476

**Authors:** Scott M. Gordon, Mailyn A. Nishiguchi, Julie M. Chase, Sneha Mani, Monica A. Mainigi, Edward M. Behrens

**Affiliations:** Division of Neonatology, Children’s Hospital of Philadelphia, Philadelphia, PA 19104, USA; Perelman School of Medicine, University of Pennsylvania, Philadelphia, PA 19104, USA; Division of Rheumatology, Children’s Hospital of Philadelphia, Philadelphia, PA 19104, USA; Center for Research on Reproduction and Women’s Health, University of Pennsylvania, Philadelphia, PA 19104, USA

## Abstract

Disruption in homeostasis of interleukin-15 (IL-15) is linked to poor maternal and fetal outcomes during pregnancy. The only cells described to respond to IL-15 at the early maternal-fetal interface have been natural killer (NK) cells. We now show a novel population of macrophages, evident in several organs but enriched in the uterus of mice and humans, expressing the β chain of the IL-15 receptor complex (CD122) and responding to IL-15. CD122+ macrophages (CD122+Macs) are morphologic, phenotypic, and transcriptomic macrophages that can derive from bone marrow monocytes. CD122+Macs develop in the uterus and placenta with kinetics that mirror interferon (IFN) activity at the maternal-fetal interface. Macrophage colony-stimulating factor (M-CSF) permits macrophages to express CD122, and IFNs are sufficient to drive expression of CD122 on macrophages. Neither Type-I nor Type-II IFNs are required to generate CD122+Macs, however. In response to IL-15, CD122+Macs activate the ERK signaling cascade and enhance production of proinflammatory cytokines after stimulation with the Toll-like receptor 9 agonist CpG. Finally, we provide evidence of human cells that phenocopy murine CD122+Macs in secretory phase endometrium during the implantation window and in first-trimester uterine decidua. Our data support a model wherein IFNs local to the maternal-fetal interface direct novel IL-15-responsive macrophages with the potential to mediate IL-15 signals critical for optimal outcomes of pregnancy.

The microarray data presented in this article have been submitted to the Gene Expression Omnibus (http://www.ncbi.nlm.nih.gov/geo/) under accession number GSE132353.

## INTRODUCTION

The notion of immune quiescence during pregnancy forms the basis for the decades-old central tenet of reproductive immunology (1). However, countless pieces of evidence now support that the action of proinflammatory cytokines are critical to a healthy gestation for both mother and fetus (2). As in many other contexts, though, the proinflammatory response must be regulated in order to avoid an adverse outcome of pregnancy caused by damage to the developing fetoplacental unit.

Perturbation in homeostasis of the pleiotropic cytokine interleukin-15 (IL-15) has been linked to poor maternal and fetal outcomes of pregnancy in mice and humans. Mice lacking IL-15 (*Il15*^*-/-*^) bear growth-restricted pups and exhibit higher rates of spontaneous resorptions than do IL-15-sufficient mice (3, 4). Mice deficient in either production of IL-15 or responsivity to IL-15 exhibit abnormal utero-placental anatomy, including impaired remodeling of the uterine spiral arteries, a pathologic hallmark of life-threatening preeclampsia in humans (5). These data translate well to findings of reduced IL-15 in human placentae of pregnancies affected by preeclampsia (6). Conversely, IL-15 permits inflammatory-mediated fetal loss in mice (4). Indeed, spontaneous and recurrent miscarriage in humans is associated with unrestrained expression of IL-15 mRNA and protein (7).

IL-15 is abundant during normal pregnancy. In mice, IL-15 mRNA is expressed in the non-pregnant uterus, as well as in the utero-placental unit throughout pregnancy, with a peak at mid-gestation (8). In humans, IL-15 mRNA is found in low abundance in proliferative phase endometrium and increases substantially in secretory phase endometrium and first trimester decidua (9). IL-15 protein has been demonstrated in endometrial stromal cells, perivascular cells abutting uterine spiral arteries, and vascular endothelial cells, echoing the presumed roles for IL-15-responsive cells in vascular remodeling.

In the current model of IL-15 signaling, IL-15 is complexed with the high-affinity α subunit of the IL-15 receptor (IL-15Rα) (10). Myeloid cells and non-hematopoietic cells then present the IL-15/IL-15Rα complex in trans to IL-15-responsive cells, typically killer lymphocytes bearing high levels of the β chain of the IL-15 receptor complex (CD122) and the common γ chain (γ_c_). Indeed, at the maternal-fetal interface, IL-15 is bound to the cell surface of CD14+ monocytes and macrophages in single cell suspensions of human first-trimester uterine decidua (9), the uterine lining remodeled to accept an embryo. Consistent with these human data, pregnant mice lacking γ_c_ but reconstituted with bone marrow from *Il15*^*-/-*^ mice exhibit normal uterine vascular remodeling. These findings support that the dominant sources of IL-15 in mice also are non-hematopoietic cells and chemo-resistant myeloid cells (11).

Altogether, these data show that appropriate signaling by IL-15 is critically important to healthy pregnancy. However, the targets and mechanisms of IL-15 signaling at the maternal-fetal interface are incompletely understood. NK cells are the prototypical IL-15-responsive cell type at the maternal-fetal interface. They are highly prevalent in the uterus and produce proinflammatory interferon γ, without which uterine arteries do not remodel and preeclampsia develops in mice (12). Because uterine and other NK cells clearly depend on IL-15 for development (3, 13), it has been presumed that NK cells are responsible for all abnormalities of pregnancy associated with disrupted homeostasis of IL-15.

We now report that novel macrophages, termed CD122+Macs, express high levels of CD122 in the uterus of mice and humans. Macrophage colony-stimulating factor (M-CSF) permits Type-I and -II interferons to drive expression of CD122 on these macrophages. CD122+Macs respond to IL-15 by activating the ERK signaling cascade, and CD122+Macs stimulated with the Toll-like receptor agonist CpG enhance production of pro-inflammatory cytokines in response to IL-15. CD122+Macs represent a new and unexpected target of IL-15 in the utero-placental unit and, thus, may have significant clinical implications in healthy and complicated pregnancies.

## METHODS

### Preparation of single-cell suspensions

Organs of interest were grossly dissected, weighed, minced finely with scissors, and digested in medium containing Liberase TM (Roche) at a final concentration of 0.28WU/mL and DNase (Roche) at a final concentration 30μg/mL for 30 min at 37°C with intermittent agitation. Cells were passed through a 70μm filter, and red blood cells were lysed with ACK lysis buffer. Cells were again filtered and counted with a Countess hemocytometer (Thermo Fisher Scientific) prior to proceeding to antibody staining for flow cytometry or cell sorting.

### Flow cytometry and cell sorting

Flow cytometry was performed on either a MacsQuant Analyzer 10 (Miltenyi), an LSRFortessa (BD), or a CytoFLEX LX (Beckman Coulter). Cell sorting was performed on either a FACSAria Fusion (BD) or a MoFlo Astrios EQ (Beckman Coulter). Data were exported as FCS files and analyzed using FlowJo 10 software. All antibody staining was performed at 4°C in the dark. Prior to fluorescent antibody staining, cells were incubated with mouse Fc block (TruStain FcX, Biolegend) or human Fc block (BD). Fixable, fluorescent LIVE/DEAD viability dye (Thermo Fisher) was used in Blue, Aqua, Green, and Near-IR per manufacturer recommendations. All antibodies, clones, and concentrations used for flow cytometry and cell sorting are listed in Supplmental Table 3.

### Microarray

Sorted cells were pelleted, removed of medium, snap-frozen on dry ice, and stored at -80°C. RNA was isolated using a Micro RNeasy mini total RNA kit (Qiagen), performed according to the manufacturer’s instructions. MouseGene 1.0ST chips were used. Microarray data were normalized by the Robust Multichip Average (RMA) algorithm and log_2_ transformed using the oligo package in R (14). Linear modeling to obtain differentially expressed genes was performed with the limma package in R (15). A gene was considered significantly differentially expressed with a FDR-adjusted p value less than 0.05. Gene ontology (GO) analysis was performed using DAVID (16, 17).

### Adoptive transfers

Single-cell suspensions of bulk bone marrow (BM) were obtained from pooled femorae, tibiae, and humeri of donor mice. Red blood cells were lysed with ACK lysis buffer, and cells were filtered through a 70μm filter. Monocytes were enriched with a mouse BM monocyte isolation kit (Miltenyi) by magnetic bead-mediated depletion of non-monocytes. Manufacturer instructions were followed exactly. Purity of transferred BM monocytes was routinely ∼90% prior to adoptive transfer.

### Bone marrow-derived macrophages (BMDMs)

Single-cell suspensions of BM were prepared as above. Adherent cells were removed by incubating bulk bone marrow cells on tissue culture-treated plates for at least 2 hours in complete DMEM (cDMEM, 10% FBS, 1% Pencillin/Streptomycin/Glutamine). Non-adherent cells were then collected, counted, and plated at 5×10^5^ cells/mL in cDMEM containing 10% L929 cell-conditioned medium (LCM, made in our laboratory) for 5-7 days. GM-BMDMs were generated by culturing non-adherent BM cells in cDMEM supplemented with 50ng/mL GM-CSF (Peprotech). CD122+BMDMs were generated by adding 1ng/mL IFNα (Biolegend) to the medium on day 5 of culture. When testing ability of IFNγ (Peprotech) to derive CD122+Macs, doses indicated were added on day 5 of culture. Anti-IFNAR and anti-IFNGR (both BioXcell) were both used at 10μg/mL in culture.

### ELISA

CD122+BMDMs were generated as above. Cells were washed on day 6 of culture, and medium was replaced with DMEM containing 3% serum for the final 16 hours of culture with or without 1μg/mL CpG 1826 (IDT), as indicated. To this medium, IL-15Rα alone (IL-15Rα Fc chimera protein, R&D Systems) or IL-15Rα pre-complexed with IL-15 (Peprotech) was added, as indicated. Final concentrations of IL-15Rα and IL-15 were 100ng/mL and 50ng/mL, respectively. Complexing of IL-15Rα with IL-15 was performed in PBS at 37°C for 30 minutes. OptEIA ELISA kits (BD) for mouse were used: IL-6 and TNF. Manufacturer instructions were followed precisely, and concentrations of cytokines were determined with a SpectraMax ELISA plate reader.

### Western blots

CD122+BMDMs were generated as above. Cells were washed on day 6 of culture, and medium was replaced with serum-free DMEM for 2 hours. Medium was replaced with cDMEM containing IL-15Rα alone or IL-15Rα/IL-15 complex, as above, for 20 minutes. Whole-cell lysates were then prepared by washing cells in cold PBS and lysing in M-PER Mammalian Protein Extraction Reagent (Thermo Scientific) with protease/phosphatase inhibitor cocktail (Cell Signaling Technology). Lysates were quantified by Bradford assay, normalized, reduced, and resolved by SDS gel electrophoresis on 10% Bis-Tris gels (Invitrogen). Resolved proteins were transferred to 0.45 μm Nitrocellulose membranes (Bio-Rad). Membranes were blocked in Odyssey Blocking Buffer (Licor) and probed with the following antibodies: pERK (Cell Signaling Technology, 1:1000), total ERK (Cell Signaling Technology, 1:1000), and GAPDH (Novus Bio, 1:2000). Membranes were incubated with the following secondary antibodies at a dilution of 1:20,000: Alexa Fluor 680 donkey anti-rabbit IgG (Invitrogen) or Alexa Fluor Plus 800 donkey anti-mouse IgG (Invitrogen). Proteins were detected and quantified using a LICOR Odyssey.

### Human endometrial samples

Endometrial biopsies were obtained from patients and volunteers at Penn Fertility Care. MAM has approval for the collection endometrial tissue (IRB# 826813). Written informed consent was obtained from all subjects. All women were between the ages of 18 and 43, with no significant medical history and regular menstrual cycles (between 25-35 days over the past 3 months). Endometrial biopsies were obtained from known fertile women 8 days after an LH surge (secretory phase), which was detected in the urine by the Clearblue Ovulation Test. All biopsies were obtained using a Pipelle Endometrial Suction Curette (Cooper Surgical). All biopsies were dissociated and analyzed within 24 hours of collection. Single cell suspensions were obtained as above.

### First trimester decidual tissue

First trimester decidual tissue was obtained from the Penn Family Planning and Pregnancy Loss center. MAM has IRB approval for the collection of first trimester extraembryonic tissue (IRB# 827072). Written informed consent was obtained from all subjects. After tissue collection, decidua was manually separated from chorionic villi based on gross morphology. All samples were dissociated and analyzed within 24 hours of collection. Single cell suspensions were obtained as above.

### Statistics

For non-microarray data, statistical analyses were performed using GraphPad Prism 8. A 1-way ANOVA with a test for linear trend was used to determine whether the slope of a line was likely to have occurred by chance alone. A (non-parametric) Wilcoxon matched-pairs signed rank test was used to compare effect of IL-15 treatment on matched samples. Mixed-effects analysis was used to compare paired observations with one-way ANOVA. Holm-Sidak’s multiple comparisons test was used in mixed-effects analyses. In all cases, *P* < 0.05 was considered significant.

### Mice

All animals were housed, cared for, and euthanized in accordance with a protocol approved by the Institutional Animal Care and Use Committee (IACUC) of Children’s Hospital of Philadelphia. Wild-type mice were strain C57Bl/6, and all knockout mice were on a C57Bl/6 background. *Ifnar*^*-/-*^, *Ifngr*^*-/-*^, and *Ifnar*^*-/-*^*Ifngr*^*-/-*^ mice, LysM-Cre and DTR-mCherry transgenic mice intercrossed to yield MM-DTR mice, and CD45.1 mice were all purchased from The Jackson Laboratory. All animals used were between 6-12 weeks of age. While it was only possible to use female mice to investigate uterine immune cells, both male and female mice were used for *in vitro* experiments, yielding similar results. For pregnant female mice, presence of copulation plugs were checked early each morning. Copulation was assumed to take place during the 12-hour dark cycle (from 6PM-6AM in our facility). Thus, E0.5 denotes the morning that a copulation plug was first detected.

## RESULTS

### CD122+ macrophages are enriched in the uterus in mice

To determine whether uterine leukocytes other than NK cells could be cellular targets of IL-15, we comprehensively examined expression of CD122 on immune cells in pregnant female mice. Populations of cells co-expressing CD122 and high levels of macrophage/monocyte-associated Fc γ Receptor 1 (FcγRI, CD64) and F4/80 were evident in several organs but were particularly enriched in the uterus of pregnant and non-pregnant mice (Fig. 1A, Supplemental Fig. 1A). Other myeloid-phenotype cells, such as Ly6G+ cells (neutrophils) and CD11c_bright_MHCII_bright_ cells (conventional dendritic cells, DCs), did not exhibit surface expression of CD122 (Fig. 1A). During gestation, these CD122+ phenotypic macrophages (herein CD122+Macs) were present throughout the maternal fetal interface: in the myometrium, mesometrial lymphoid aggregate of pregnancy (MLAP), decidua, and placenta (Fig. 1B).

**Figure 1.**
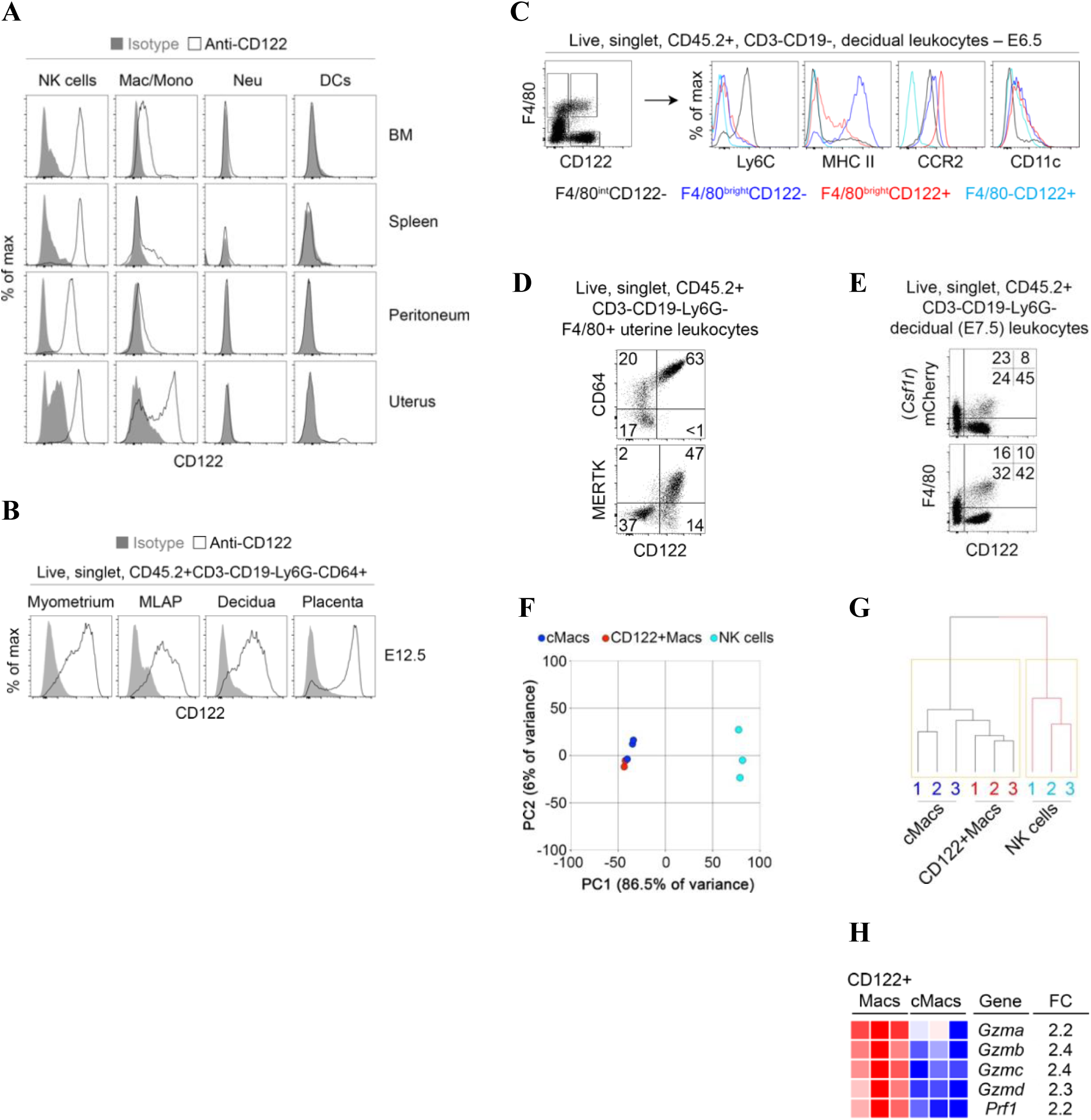
Macrophages expressing CD122 are evident in several organs but are enriched in the uterus. (A) Strong expression of CD122 is evident on macrophage-lineage cells but not other myeloid cells in the bone marrow and uterus. Shaded grey histograms represent isotype control antibody staining, while open histograms show staininig with anti-CD122. Indicated populations are gated on live, singlet, CD45.2+ CD3-CD19- leukocytes in non-pregnant mice. “NK cells” are Ly6G-CD64-NKp46+, macrophages and monocytes (“Mac/Mono”) are Ly6G-CD64+NKp46-, neutrophils (“Neu”) are Ly6G+CD64-, and dendritic cells (“DCs”) are Ly6G-CD64-NKp46-CD11c+. Data are representative of at least 10 mice over at least 3 independent experiments. (B) At midgestation in pregnant mice, CD122+Macs are evident in all layers of the maternal-fetal interface. Data are representative of 3 independent experiments with 2-3 mice per experiment. (C) Phenotypically, CD122+F4/80_bright_ cells are macrophage-like but express less MHC Class II and more CCR2 than CD122-F4/80_bright_ cells. Data are representative of at least 5 independent experiments with 2-3 mice per experiment. (D) Uterine CD122+Macs from non-pregnant mice are largely CD64+ and MERTK+. Data are representative of at least 2 independent experiments with 4 total mice. (E) Co-expression of *Lyz2* (LysM) and *Csf1r* (M-CSFR) in CD122+Macs. In “MM-DTR” mice, LysM-Cre acts on a transgene containing *Csf1r* promoter elements, followed by a floxed STOP cassette, followed by an mCherry-Diphtheria toxin receptor fusion protein to specifically label macrophages. Data are representative of at least 5 independent experiments, with at least 2 mice per experiment. (F, G) Transcriptionally, CD122+Macs cluster with CD122-conventional macrophages (cMacs) by (F) principal component analysis (PCA) and (G) hierarchical clustering analysis (HCA). Macs cluster away from NK cells (CD122+F4/80-). Prior to PCA and HCA, control probes and low-variance probes in microarray data were filtered out. Indicated cells were FACS-sorted from 3 independent groups of pooled E7.5 deciduae, with each group consisting of 4-5 mice. (H) Heat map depicting CD122+Macs are enriched in cytolytic transcripts relative to cMacs.

Rarely are classical myeloid and lymphoid proteins co-expressed in the same cell. Recently, NK cells expressing canonical myeloid transcripts, including *Csf1r*, were identified in obese mice and humans (18). Thus, it was possible that CD122+Macs represented a subpopulation of so-called myeloid NK cells, given the high abundance of NK cells in the early gestation uterus. In order to determine to which lineage uterine CD122+Macs cells belonged, we compared their morphology to other uterine cell types. Murine uterine CD122+F4/80_bright_ cells were morphologically large and hyper-vacuolated, similar to conventional CD122-macrophages (cMacs, Supplemental Fig. 1B, 1C). The cell surface phenotype of CD122+Macs overlapped substantially with that of cMacs. Most CD122+Macs and cMacs were CD11c_int_, Ly6C_low/neg_, CCR2_hi_, and MERTK_hi_, a constellation of findings typically associated with macrophages (Fig. 1C, 1D). Aside from CD122, CD122+Macs and cMacs could be distinguished by expression of MHC Class II (MHCII), with CD122+Macs largely MHCII_low_ and cMacs largely MHCII_hi_ (Fig. 1C).

In contrast to CD122+Macs and cMacs, the morphology and flow cytometric profile of CD122+F4/80-CD64-cells were consistent with classical large, granular, decidual NK cells (Fig. 1C, Supplemental Fig. 1B-E). Nearly all decidual NK cells express NKp46 and the tissue-resident NK cell marker CD49a, especially early in gestation (19). We found that CD122+Macs did not express CD49a, CD49b/DX5, T-bet or Eomesodermin (Supplemental Fig. 1E), closely-related master regulators of innate lymphoid fate that are abundantly expressed in uterine innate lymphoid cells (19-21). Morphologic and phenotypic data thus supported the notion that CD122+Macs cells are indeed macrophage-lineage cells and not NK cells, DCs, or neutrophils.

We next tested whether CD122+Macs were most like macrophages or another cell type at the level of the transcriptome. A monocyte-macrophage-specific reporter system, known as the MM-DTR mouse, fluorescently labels with mCherry monocytes and macrophages that transcribe both *Lyz2* (LysM) and *Csf1r* (macrophage colony-stimulating factor receptor, M-CSFR) (22). Expression of CD122 directly correlated with expression of mCherry in F4/80_bright_ cells in the uterus (Fig. 1E). We then sort-purified decidual CD122+Macs, cMacs, and NK cells to a purity of at least 95% (Supplemental Fig. 1F) and analyzed the transcriptome using microarrays. We found that 86.5% of the variance in gene expression among these populations could be attributed to the first principal component, PC1 (Fig. 1F). Clustering samples along PC1 clearly separated cMacs and CD122+Macs from NK cells. Similarly, hierarchical clustering analysis showed that all macrophages were closely related to each other but distantly related to NK cells (Fig. 1G). Among genes enriched in CD122+Macs relative to NK cells were those classically associated with macrophages, including numerous complement receptors, Toll-like receptors, and *Cd68, Csf1r, Fcgr1, Lyz2*, and *Cx3cr1* (Supplemental Table 1). Among genes enriched in NK cells relative to CD122+Macs were those typically associated with innate and killer lymphocytes, including *Ncr1* (NKp46), transcripts encoding cytolytic mediators, Ly49 receptors, and *Eomes* (Eomesodermin) and *Tbx21* (T-bet). Of note, some cytolytic genes, including *Prf1* and several granzymes, were modestly but significantly enriched in CD122+Macs relative to cMacs (Fig. 1H, Supplemental Table 2). Altogether, these data support the notion that CD122+Macs and cMacs are highly related but distinct subtypes of uterine macrophages.

### CD122+Macs are present in human endometrium and decidua

Given the unusual phenotype and restricted anatomic location of CD122+Macs in mice, we assessed whether such macrophages were also found in the human uterus. We therefore examined human secretory endometrium during the implantation window, as well as human first-trimester decidua, for the presence of human CD122+Macs (hCD122+Macs). We relied on expression of CD14 and CD64 to identify human classical monocytes and macrophages by flow cytometry (Supplemental Fig. 3A). We defined human NK (hNK) cells as CD14-CD56_bright_ (uterine) or CD14-CD56_dim_ (blood). As expected, hNK cells were abundant in the uterus and expressed high levels of CD122.

We then identified CD14/CD64+ cells enriched in the human pre- and post-implantation uterus that expressed CD122 (Fig. 2A-C, Supplemental Fig. 2A-E, Supplemental Fig. 3A-C). These hCD122+Macs varied widely in abundance among individual samples but were present in all first-trimester deciduae tested (Fig. 2A-C, Supplemental Fig. 2A-E). Human CD122+Macs were also present in over half of human secretory phase endometrial biopsies examined during the implantation window (Fig. 2C, Supplemental Fig. 3A-C). We found that hCD122+Macs often expressed CD16 and variably expressed CD11c, as did human conventional macrophages (hcMacs, Fig. 2B and 2C, Supplemental Fig. 2A-E, Supplemental Fig. 3A). Both hCD122+Macs and hcMacs expressed HLA-DR, but in some samples, hCD122+Macs expressed modestly less HLA-DR, similar to our observations of MHCII levels on CD122+Macs and cMacs in the mouse.

**Figure 2.**
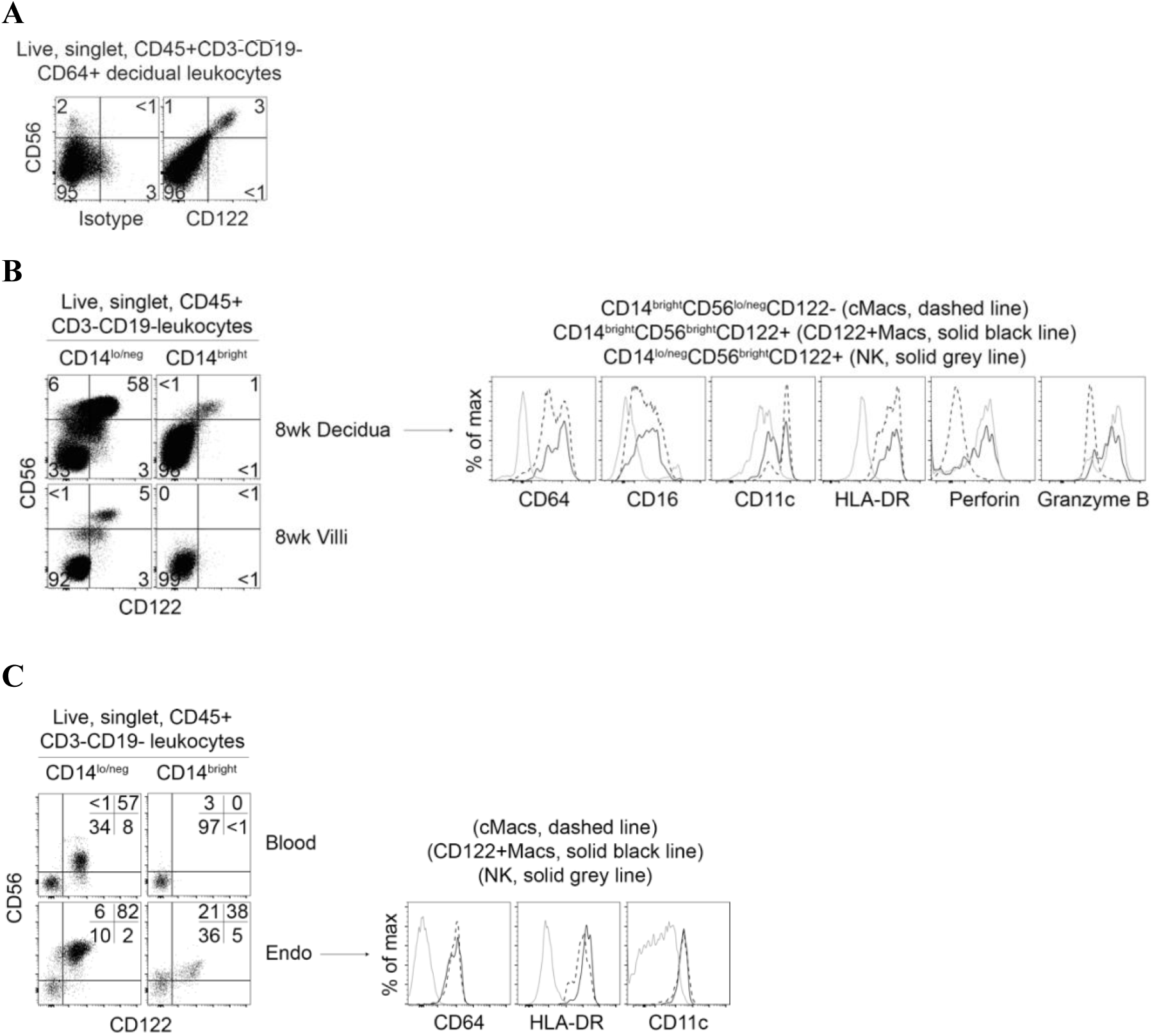
Human CD122+Macs are evident in the decidua before and during pregnancy. (A) Human equivalent of murine CD122+Macs in first-trimester decidual tissue from 6-week elective termination of pregnancy, stained with isotype control (left panel) or anti-CD122 (right panel). Data are representative of 3 independent experiments. (B) Human CD122+Macs in first-trimester decidual tissue, but not first-trimester chorionic villi, from 8-week elective termination of pregnancy. Phenotypically, human CD122+Macs express many markers of the macrophage lineage. Human CD122+Macs diverge from human cMacs by expressing high levels of CD122, CD56, Perforin, and Granzyme B. Additional examples are shown in Supplemental Figure 2. (C) Human CD122+Macs in secretory phase endometrium during the implantation window but not in blood from the same subject. Additional examples are shown in Supplemental Figure 3.

Human CD122+Macs variably expressed CD56 (Fig. 2A-C, Supplemental Fig. 2A-E, Supplemental Fig. 3A-C). CD56 is typically associated with cytotoxic lymphocytes, but it has been found on myeloid cells in certain contexts (23). Consistent with this literature, human CD122+Macs expressing higher levels of CD56 appeared to express the cytolytic molecules Perforin and Granzyme B at the protein level (Fig. 2B, Supplemental Fig. 2B, 2D and 2E, Supplemental Fig. 3A). These data agree with our transcript-level data in mice, showing enrichment of cytolytic mediators in CD122+Macs relative to cMacs (Fig. 1H, Supplemental Table 2). Overall, human CD122+Macs bore striking resemblance to murine CD122+Macs, supporting the use of our murine model to obtain mechanistic insights into the biology of this novel uterine macrophage in humans.

### Mouse CD122+Macs can derive from monocytes

Our morphologic, phenotypic, and transcriptomic data supported that CD122+Macs were macrophages. Myriad embryonic and adult progenitors give rise to macrophages in different tissues (24, 25). Monocytes are recruited in large numbers to the gravid uterus during gestation (26). We thus tested the hypothesis that CD122+Macs could derive from adult bone marrow monocytes. Magnetic bead-purified Ly6C_hi_ monocytes from adult mouse bone marrow of MM-DTR mice were adoptively transferred into pregnant recipients during the peri-implantation period (Fig. 3A and 3B). Recipients were then sacrificed at mid-gestation. We observed tissue-specific differences in phenotype of recovered donor cells with respect to both F4/80 and CD122. Donor cells recovered from recipient blood, spleen, and bone marrow were almost uniformly F4/80_int_ (Fig. 3B), suggesting maintenance of monocyte fate. In contrast, donor cells recovered from recipient peritoneum, myometrium, and decidua contained F4/80_bright_ cells, suggesting some conversion to tissue macrophages in these organs.

**Figure 3.**
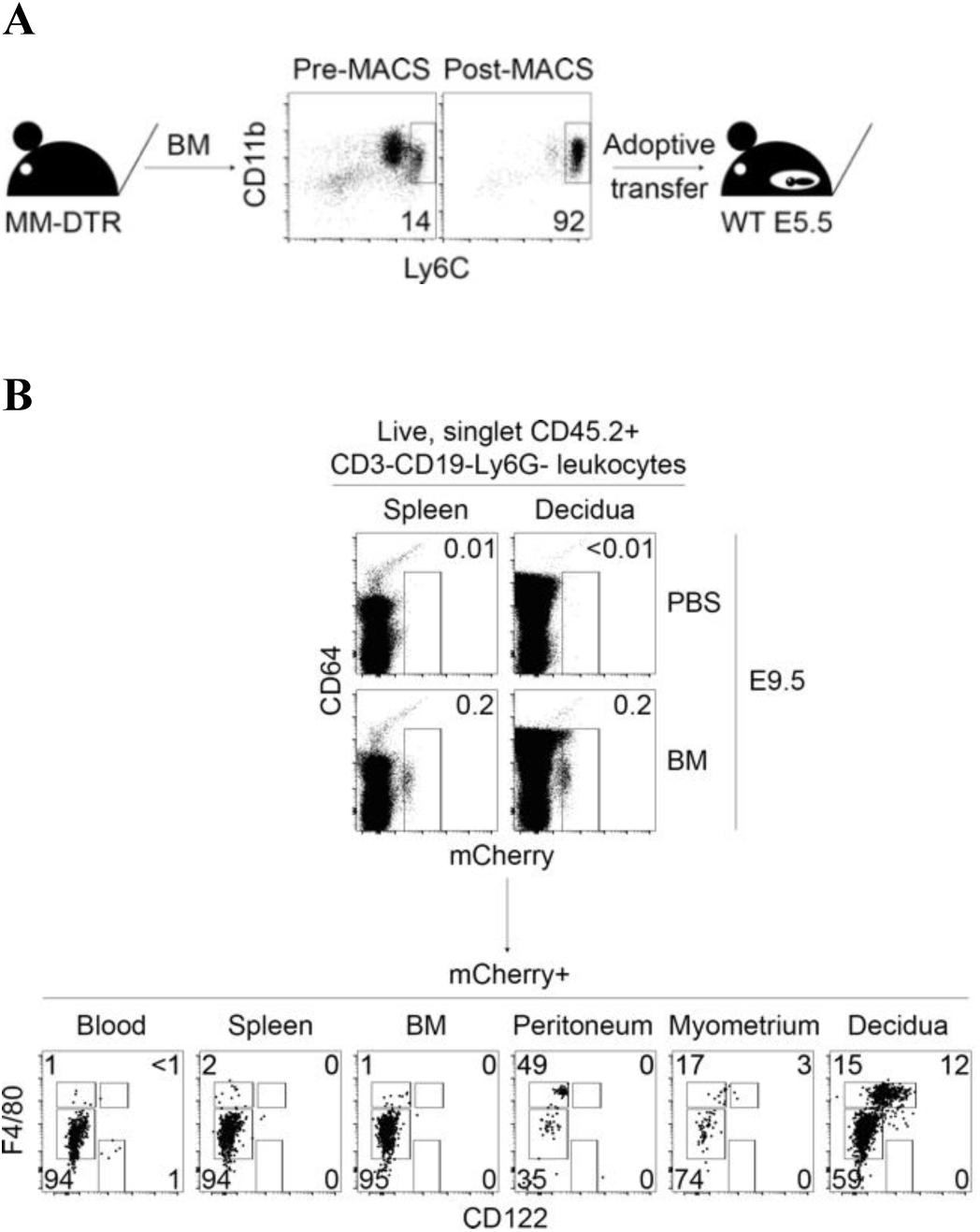
Bone marrow monocytes can give rise to CD122+Macs at the maternal-fetal interface during pregnancy. (A) Bone marrow monocytes from MM-DTR mice were enriched by magnetic-activated cell sorting (MACS) from non-pregnant mice prior to adoptive transfer into post-implantation pregnant mice. (B) Uterine CD122+Macs can derive from bone marrow (BM) monocytes. (Top panels) Four days after i.v. adoptive transfer of MM-DTR BM monocytes, mCherry+ (donor-derived) cells were recovered from recipients. Control mice received i.v. saline (PBS). Data are representative of at least 3 independent experiments with 2-3 recipients per experiment.

Despite conversion of monocytes to F4/80_bright_ cells in the peritoneum, we found that transferred monocytes converted to macrophages expressing high levels of CD122 only in the uterus (Fig. 3B). These data are consistent with our findings that CD122+Macs are enriched in the uterus in the steady state. Of note, we performed this experiment with both pregnant and non-pregnant donors with identical results (data not shown), suggesting no intrinsic differences in potential of bone marrow monocytes between pregnant and non-pregnant mice. Altogether, our data support a model in which bone marrow monocytes are recruited to the uterus and differentiate into CD122+Macs.

### Tissue-specific population dynamics of CD122+Macs during gestation

Populations of myeloid cells, including dendritic cells, monocytes, and macrophages, are in constant flux at the maternal-fetal interface during gestation (26, 27). We thus investigated population dynamics of uterine CD122+Macs before and during pregnancy in the mouse. The myometrium exhibitied progressive percent enrichment of F4/80_bright_ cells over the course of gestation (Fig. 4A and 4B), which agrees with and extends prior findings (26). This was in contrast to progressive declines in NK cells after mid-gestation (Fig. 4A-C), consistent with prior reports (28). On a percentage basis, myometrial CD122+Macs were more apparent over the course of gestation until E16.5 (Fig. 4A and B). By E12.5 and before E16.5, myometrial CD122+Macs also exhibited more robust expression of CD122 on a per-cell basis. The absolute numbers of CD122+Macs per gram of myometrial tissue remained relatively stable through mid-gestation but declined thereafter (Fig. 4C).

**Figure 4.**
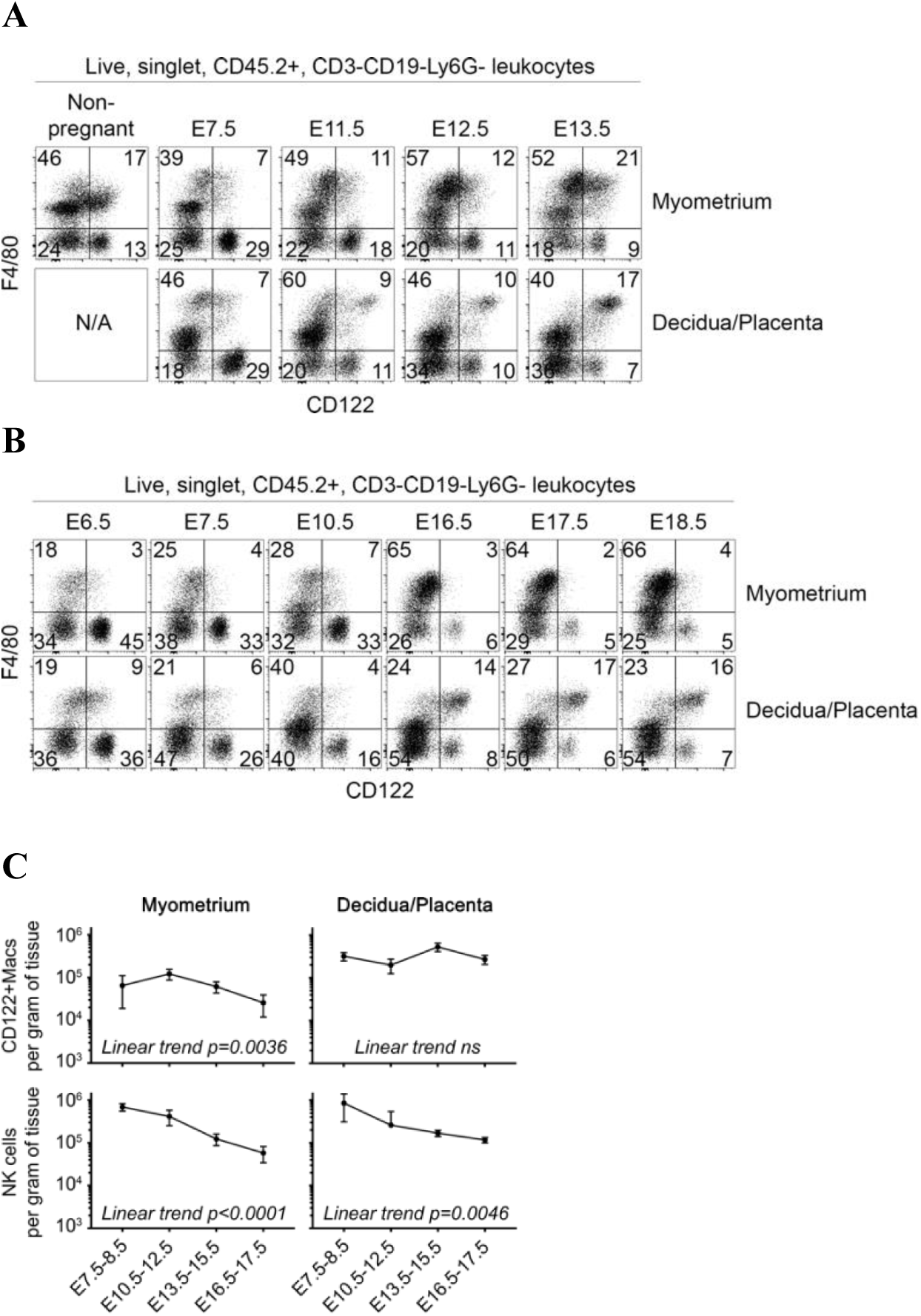
Population dynamics of CD122+Macs differ between myometrium and deciduo-placental unit. (A, B) The myometrium is progressively enriched in CD122+Macs over the course of gestation, until the sudden decline of CD122+Macs at E16.5 and beyond. The decidua or combined deciduo-placental unit is progressively enriched in CD122+Macs as gestation progresses toward term. “N/A”, not applicable. Data are representative of at least 3 independent experiments with at least 3 total mice per timepoint. (C) Quantification of numbers of CD122+Macs and NK cells per gram of indicated tissue shows that CD122+Macs decline over gestation in the myometrium but remain constant in the decidua/placenta. NK cells decline over time in both the myometrium and decidua/placenta. P values were generated by one-way ANOVA with a test for linear trend and refers to whether the slope of each line (showing changes in cell number per gram of tissue over time) was likely to have occurred by chance alone. “ns”, not significant, p=0.38.

Similar to the myometrium, the combined decidual and placental layers exhibited progressive decline in NK cells but progressive relative enrichment of CD122+Macs during pregnancy (Fig. 4A and 4B). Unlike the myometrium, however, this relative enrichment of CD122+Macs in the decidua and placenta appeared to be at the expense of F4/80_bright_ cMacs in those tissue layers. Also unlike the myometrium, a robust proportion and absolute number of CD122+Macs persisted as gestation approached term (Fig. 4B and 4C). In the early post-implantation period through E10.5, CD122+Macs expressed moderate amounts of CD122 in the decidua and placenta (Fig. 4A and 4B). From E11.5 through E18.5, the last timepoint we examined before delivery, deciduo-placental CD122+Macs exhibited progressively more robust expression of CD122 on a per-cell basis.

In the non-pregnant uterus, we observed a population of CD122+F4/80_int_ cells that became less apparent but persisted throughout pregnancy (Fig. 4A and 4B). These cells could have represented monocytes/macrophages or F4/80+ uterine eosinophils (29). Our data confirmed that presumptive eosinophils, with high side scatter properties and low to no expression of CD64, are abundant within the bulk F4/80_int_ population (data not shown). However, these cells were uniformly negative for CD122. F4/80_int_CD122+ cells, on the other hand, exhibited lower side scatter properties and were CD64+, consistent with monocytes/macrophages. These data reinforce a model in which monocytes and macrophages are the only uterine myeloid cells that express surface CD122. Further, these data provide evidence for a model in which populations of CD122+ monocytes and macrophages are dynamically regulated in a tissue layer-specific fashion in the pre-pregnant and gravid uterus.

### Type-I and II interferons are sufficient for development of CD122+Macs

Our data suggested that the uterus strongly favored development of CD122+Macs. Further, the drivers of the CD122+Mac fate appeared transiently strongest just after mid-gestation in the myometrium and persistently strong from mid-gestation through near-term in the deciduo-placental unit. Total interferon (IFN) activity at the mouse maternal-fetal interface precisely mirrors the population dynamics of CD122+Macs (30). The non-pregnant uterus transcribes Type-I IFN and IFN-stimulated genes (ISGs) (31), and Type-II IFN is apparent during the estrous phase (32). In the pregnant myometrium, total IFN activity is low until E10, peaks between E11 and E15, then returns to low levels after E15 (30). IFN activity is already robust in the developing placenta at E10 but spikes dramatically after E10 through to term.

To understand whether IFNs played a role in development of uterine CD122+Macs, we first compared the transcriptomes of decidual cMacs and CD122+Macs during early gestation, when both cell types were present and in similar abundance. Review of individual transcripts and gene ontology (GO) analysis of transcripts significantly upregulated in CD122+Macs relative to cMacs showed strong enrichment of genes associated with interferon signaling and antiviral responses (Fig. 5A, Supplemental Table 2). With this information, we next sought to determine whether IFNs were sufficient to drive expression of CD122 on decidual macrophages. F4/80+ cells in bulk single-cell suspensions of early gestation decidual capsules exhibited a dose-dependent upregulation of surface CD122 upon exposure to recombinant IFNα, though this effect was strongest in F4/80_int_ cells (Fig. 5B and data not shown). To extend these findings, we cultured sort-purified uterine F4/80_int_ monocytes in macrophage colony-stimulating factor (M-CSF) with or without Type-I IFNα. Consistent with the notion that culture in macrophage colony-stimulating factor (M-CSF) results in endogenous production of Type-I IFN by macrophages (33), we did observe some expression of CD122 on uterine monocytes cultured in M-CSF alone (Fig. 5B and 5C). However, addition of IFNα resulted in robust upregulation of CD122+ compared to culture in M-CSF alone.

**Figure 5.**
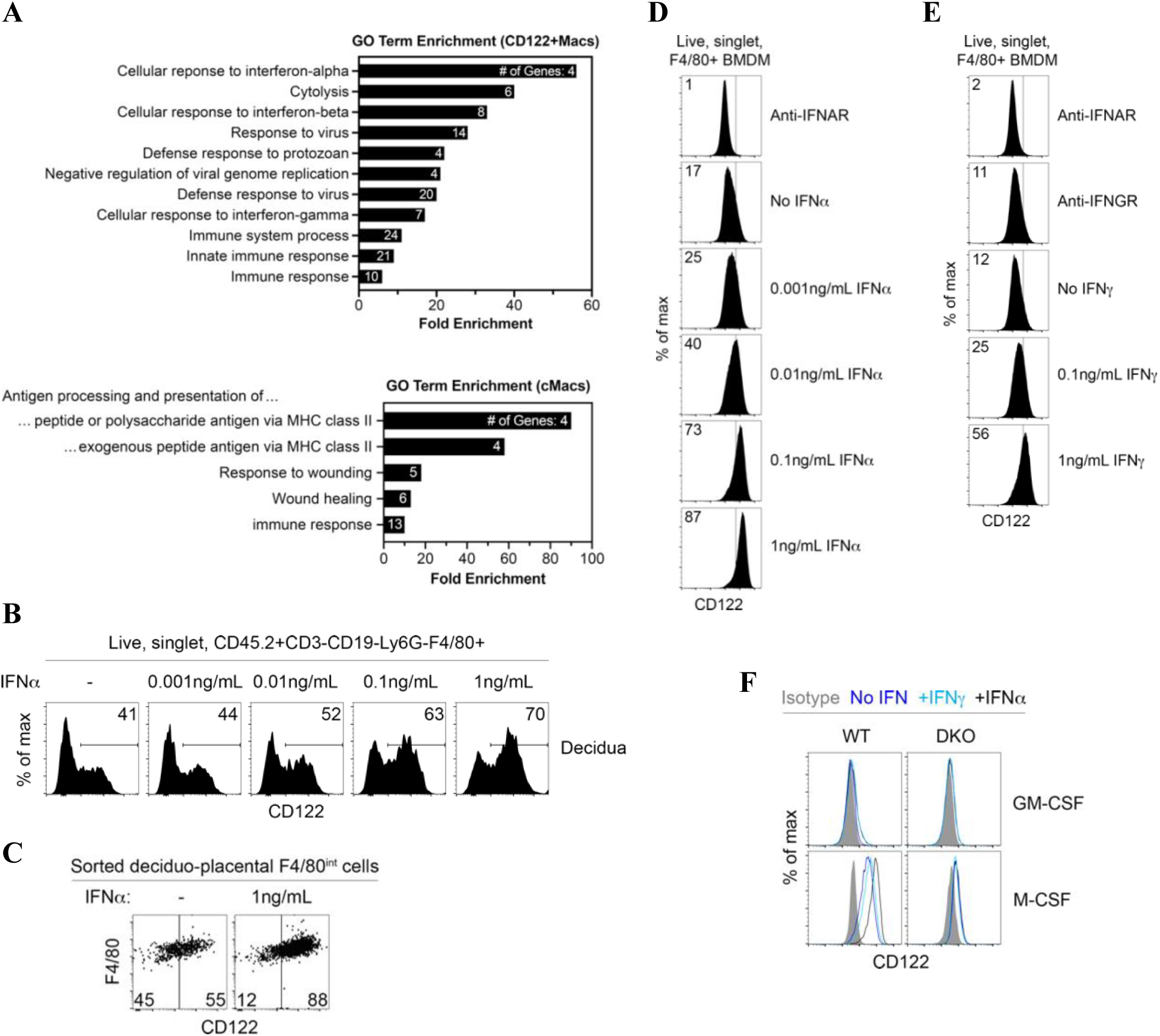
M-CSF and interferons drive expression of CD122 on uterine and bone marrow-derived macrophages. (A) Interferon-responsive genes (ISGs) are overrepresented among genes significantly enriched in E7.5 decidual CD122+Macs relative to cMacs by microarray. Gene ontology (GO) analysis by DAVID of all genes significantly enriched in CD122+Macs relative to cMacs. Bars show fold enrichment for indicated GO Biological Process (GO BP) terms significantly enriched with FDR-adjusted p<0.05. GO terms are represented by the number of genes shown in each bar. (B) Decidual F4/80+ cells upregulate CD122 in a dose-dependent fashion after 18hrs in culture with M-CSF and IFNα. Bulk E6.5 decidual cell suspensions were cultured in indicated concentrations of IFNα. Data are representative of 2 independent experiments. (C) Monocytes (CD122-F4/80_int_) sort-purified from E15.5 decidua/placenta upregulate CD122 after 24 hours in culture with M-CSF and IFNα. Data are representative of 2 independent experiments with 2-5 mice per experiment. (D, E) Dose-dependent upregulation of CD122 on bone marrow-derived macrophages (BMDMs) cultured with M-CSF and (D) Type-I IFNα or (E) Type-II IFNγ. Data are representative of at least 5 independent experiments. (F) Culture of bone marrow in M-CSF, not granulocyte/macrophage colony-stimulating factor (GM-CSF), permits induction of CD122 in response to IFN. M-CSF also drives modest expression of CD122 independent of IFN. “DKO” denotes IFNAR/IFNGR double KO mice. Data are representative of 3 independent experiments with 2-3 mice per group.

Further, we could derive CD122+Macs *in vitro* from nonadherent bone marrow cells. Traditional bone marrow-derived macrophages cultured with M-CSF (BMDMs) upregulated CD122 in a dose-dependent fashion in response to both Type-I IFNα and Type-II IFNγ (Fig. 5D and 5E). While the ability to drive expression of CD122 on BMDMs was shared by both Type-I and II IFNs, we did observe differential effects of Type-I and -II IFNs on expression of MHCII (Supplemental Fig. 4A), in agreement with prior reports (34). Consistent with endogenous production of Type-I IFN by M-CSF-stimulated macrophages (33), blockade of the Type-I IFN receptor (IFNAR) nearly abolished expression of CD122 by BMDMs cultured in M-CSF alone (Fig. 5D and 5E). Blockade of the IFNγ receptor (IFNGR), however, had no effect on level of CD122 in BMDMs cultured in M-CSF alone, suggesting that BMDMs do not produce endogenous IFNγ. Use of combined Type-I/Type-II IFN receptor knockout BMDMs (*Ifnar-/-Ifngr-/-* double-knockout, DKO) showed that DKO BMDMs appropriately do not upregulate CD122 with IFN treatment (Fig. 5F). However, we detected modest but nonzero levels of CD122 on DKO BMDMs by virtue of culture in M-CSF. Thus, IFNs enhance expression of CD122, but they are not strictly required for expression of CD122 on BMDMs.

To determine whether expression of CD122 by BMDMs in response to IFNs was a phenomenon universal to all macrophages, we generated “GM-BMDMs” by culturing nonadherent bone marrow in medium supplemented with high-dose granulocyte-macrophage colony-stimulating factor (GM-CSF), instead of M-CSF (35, 36). Consistent with prior reports (35), GM-BMDMs upregulated CD11c and MHCII to a greater extent than traditional BMDMs (Supplemental Fig. 4B). Also in contrast to traditional BMDMs, GM-BMDMs produce less endogenous Type-I IFN (36). Indeed, we observed that GM-BMDMs exhibited little to no expression of CD122 relative to traditional BMDMs (Fig. 5F, Supplemental Fig. 4B). While traditional BMDMs upregulated CD122 robustly in response to Type-I IFN, GM-BMDMs did not change expression of CD122 in response to exogenous Type-I IFN. Taken together, these data support a model in which M-CSF primes bone marrow monocytes to develop into macrophages capable of upregulating CD122 in response to IFNs.

### Type-I and II interferons are not required for development of CD122+Macs in vivo

We next assessed whether responsivity to Type-I and/or II IFNs was required for development of uterine CD122+Macs *in vivo*. As CD122+Macs could derive from monocytes, we chose to first determine requirements for Type-I and II IFNs in generation of CD122+Macs with adoptive transfer of BM monocytes. While recovery of adoptively transferred cells from mice not treated with radiation or chemotherapy is low, this approach allows for testing of cell-intrinsic requirements for Type-I and -II IFNs in development of uterine CD122+Macs in normal pregnancy. As discussed above, both Type-I and -II IFNs are produced at the maternal-fetal interface (30-32). Recently, Type-III IFNλ was shown to play a critical role in fetal protection against infection with Zika virus in mice and humans (37, 38). Thus, all three known families of IFN are actively produced at the maternal-fetal interface. Combined with our data that either Type-I or -II IFN is sufficient to induce expression of CD122 on BMDMs in vitro, we hypothesized that there would be no specific IFN required for development of CD122+Macs. In other words, any class of IFN might be able to signal to uterine macrophages to differentiate into CD122+Macs. Consistent with this hypothesis, donor-derived CD122+Macs were equally evident after transfer of BM monocytes from wild-type, Type-I IFN receptor-deficient (IFNAR KO), Type-II IFN receptor-deficient (IFNGR KO), and DKO mice (Fig. 6A and 6B). Consistent with these data, we also found similar abundance of uterine CD122+Macs in non-pregnant IFNAR KO, IFNGR KO, and DKO mice (Fig. 6C). Altogether, these data support that receptivity to Type-I and -II IFNs, alone or in combination, is not required for BM monocytes to reach the uterus and develop into CD122+Macs.

**Figure 6.**
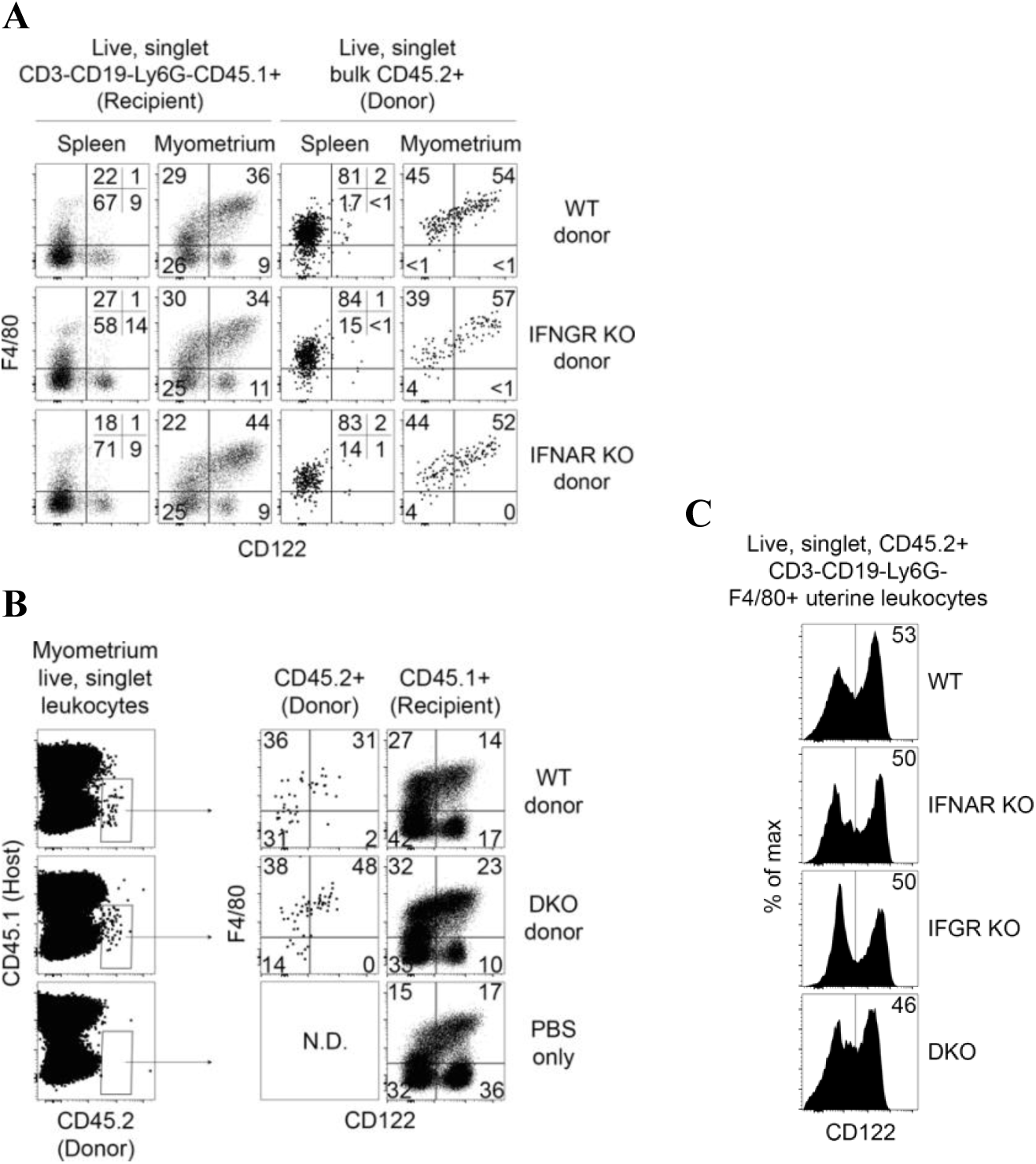
Neither Type I nor Type II IFN, alone or in combination, is required to generate uterine CD122+Macs *in vivo*. (A) MACS-enriched BM monocytes singly-deficient in either Type-I IFN (IFNAR KO) or Type-II IFN receptor (IFNGR KO) are capable of giving rise to uterine CD122+Macs. (B) MACS-enriched BM monocytes doubly-deficient in both IFNAR and IFNGR KO (double KO, DKO) are capable of giving rise to uterine CD122+Macs. BM monocytes from indicated non-pregnant donor mice (CD45.2+) were adoptively transferred into congenic (CD45.1+) peri-implantation pregnant recipient mice. Indicated organs were analyzed at E15.5. Late in gestation, transferred cells were recovered predominantly from myometrium and not from decidua/placenta. Data are representative of 2-3 recipients from at least 2 independent experiments. (C) Development of uterine CD122+Macs in vivo in non-pregnant wild-type, IFNAR KO, IFNGR KO, and DKO mice. Data are representative of 4-6 total mice over 2 independent experiments.

### IL-15 signals through CD122 to enhance function of CD122+Macs

Despite the observed expression of CD122 by CD122+Macs, if and how IL-15 signals to macrophages remained unknown. IL-15 has been shown to signal through CD122 and the common gamma chain (γ_c_), which results in phosphorylation of numerous downstream targets, including ERK (39). To test for phosphorylation of ERK in CD122+Macs exposed to IL-15, we first created CD122+BMDMs by culturing nonadherent bone marrow cells in M-CSF plus IFNα. Cytokines already present in culture, including exogenous IFNα, were then removed by washing adherent cells thoroughly and adding fresh, cytokine-free medium. Washed CD122+BMDMs were then stimulated in the presence or absence of IL-15 pre-complexed to the α chain of the IL-15 receptor (IL-15Rα), which most closely approximates true presentation of IL-15 *in vivo* and optimizes its activity (40). BMDMs stimulated with IFN express IL-15Rα (41), which can transduce IL-15 signals in the absence of CD122 in certain cell types (42). Use of a pre-associated IL-15/IL-15Rα complex, rather than free IL-15, ensured that any effects seen by stimulating CD122+BMDMs with IL-15 would be mediated through CD122/γ_c_ and not IL-15Rα. After 30 minutes of stimulation, we observed robust phosphorylation of ERK1/2 in cells treated with the IL-15 complex (Fig. 7A and 7B). These data support that IL-15 signals through CD122 in macrophages using the ERK/MAPK cascade, similar to other IL-15-responsive cell types.

**Figure 7.**
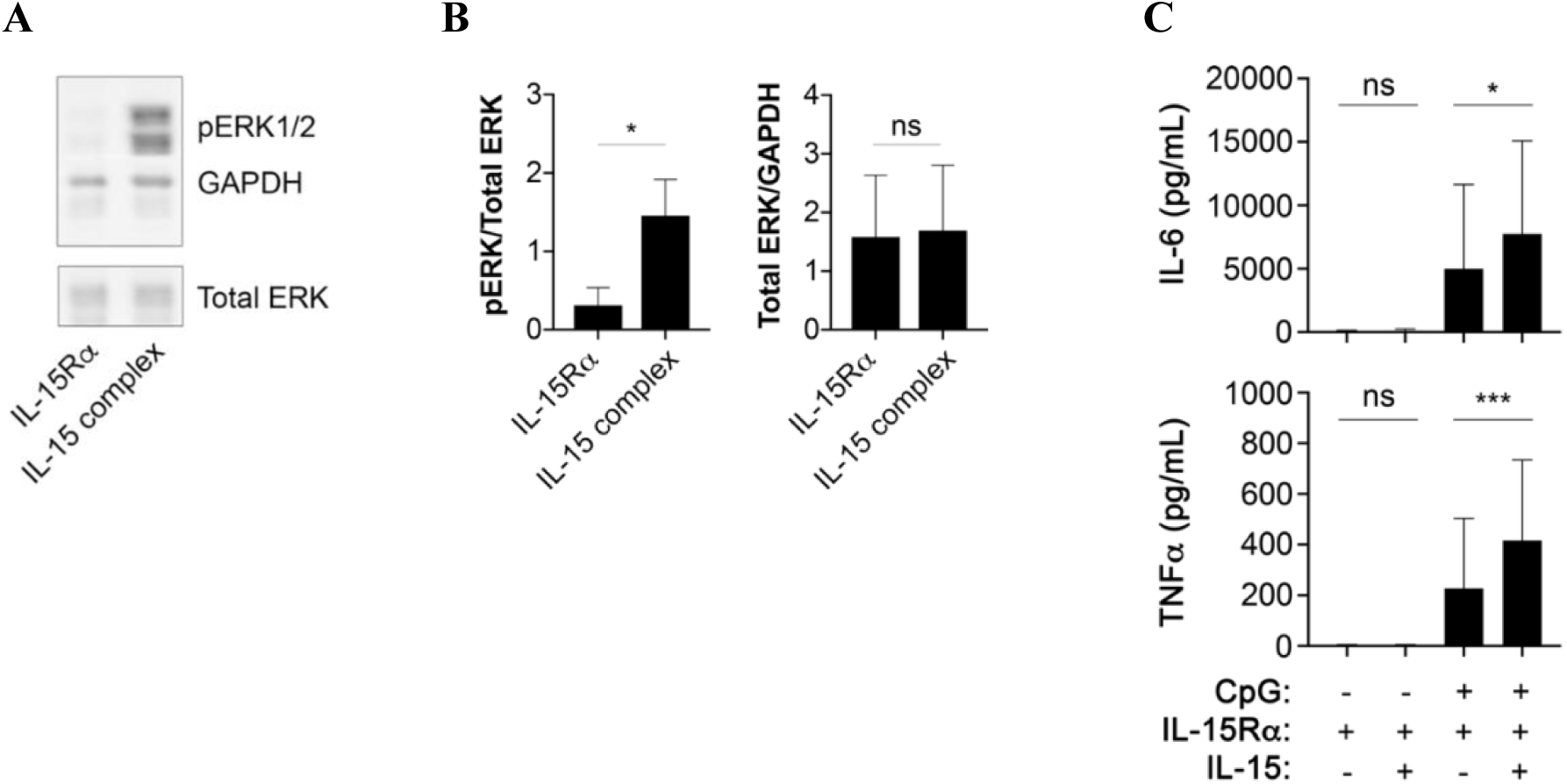
CD122+Macs signal and exhibit enhanced function in response to IL-15. (A) Phosphorylation of ERK1/2 (pERK1/2) by Western blot of CD122+BMDMs after 30 minutes of stimulation with IL-15 complex (recombinant murine IL-15 pre-complexed to recombinant murine IL-15Rα). “IL-15Rα” denotes addition of only recombinant IL-15Rα. (B) Increased pERK by Western is not due to increased total ERK. *, p=0.016 by the Wilcoxon matched-pairs signed rank test, includes 7 individual mice over 2 independent experiments. “ns”, not significant. (C) CD122+BMDMs co-stimulated with IL-15 complex and CpG produce more TNFα and IL-6 by ELISA than CD122+BMDMs stimulated with CpG alone. Here, presence of both IL-15Rα and IL-15 denotes use of IL-15 complex. BMDMs were stimulated for 16-24 hours, after which supernatants were collected for ELISA. *p=0.015 (top panel), ***p=0.0003 (bottom panel) by mixed-effects analysis (one-way ANOVA with paired observations) with Holm-Sidak’s multiple comparisons test. Bar graphs are compiled data from 11 biological replicates from 5 independent experiments, with BMDMs in triplicate generated from 1-3 individual mice per experiment. All individual data points, showing consistent response of BMDMs to IL-15 but variable response to CpG, are plotted in Supplemental Fig. 5.

We next tested whether IL-15 could act on CD122+Macs to modulate production of cytokines. CD122+BMDMs stimulated with the Toll-like receptor 9 (TLR9) agonist CpG produced abundant tumor necrosis factor α (TNFα) and IL-6 (Fig. 7C). Those CD122+BMDMs co-stimulated with CpG and IL-15 pre-complexed to IL-15Rα exhibited enhanced production of TNFα and IL-6 (Fig. 7C, Supplemental Fig. 7), but we did not observe changes in levels of IL-10 (data not shown). Overall, these data support that CD122+Macs are novel cellular targets for IL-15.

## DISCUSSION

Macrophages are critical for successful pregnancy. Severe abnormalities of pregnancy are found in osteopetrotic (*op/op*) mice that lack functional M-CSF and exhibit substantially reduced uterine macrophages (43). Implantation of the embryo is compromised and litter sizes are small relative to M-CSF-sufficient mice. *Op/op* dams cannot be mated with *op/op* sires, as the complete absence of M-CSF is incompatible with gestation of live litters. These data complement a later study showing that inducible deletion of macrophages early in pregnancy causes complete fetal loss, because macrophages sustain the vasculature of ovarian corpus luteum, required to produce progesterone and maintain early pregnancy (29).

Macrophages are abundant at the maternal-fetal interface and have been shown to directly abut blood vessels and NK cells (44). Fetal trophoblasts of the developing placenta are thought to communicate bidirectionally with macrophages, but few studies have formally addressed this (45, 46). These findings suggest that uterine and placental macrophages serve numerous critical functions in pregnancy but are likely heterogenous. Several groups have investigated macrophage heterogeneity during pregnancy in mice and humans (47, 48), but subsets of uterine macrophages remain incompletely described. In this work, we address a critical unmet need to gain a deeper understanding of signals that drive phenotype and function of macrophages at the maternal-fetal interface.

We found novel and unexpected uterine macrophages that express CD122 and respond to IL-15 under direction of M-CSF and IFNs. While IL-2 also signals to cells expressing CD122 and γ_c_, IL-2 is absent from the maternal-fetal interface in the steady state (8). These data suggest that CD122+Macs respond specifically to IL-15 during steady state pregnancy. Gene ontology analysis confirmed that the transcriptome of CD122+Macs is enriched in ISGs. IFNs are classically produced in response to viral infections, but IFNs are abundant in the cycling uterus and at the maternal-fetal interface in the steady state. Type-I and -II IFNs were sufficient to drive expression of CD122 on uterine and bone marrow-derived monocytes and macrophages. However, neither Type-I nor Type-II IFN was required, alone or in combination, to generate CD122+Macs in the uterus of knockout mice or from BM monocytes adoptively transferred into pregnant hosts. Further, M-CSF alone could drive modest expression of CD122 on cultured BMDMs independently of IFN. These data suggest that a combination of factors, likely including some we have yet to investigate, promote development of CD122+Macs in the uterus. We also acknowledge the possibility that Type-III IFN could play a role in shaping the fate of CD122+Macs, though this remains to be tested. Type-III IFN is the most recently described IFN, and extremely limited information is available about its presence at the maternal-fetal interface (38). Like Type-I and -II IFN, Type-III IFN signals through STAT1 and activates networks of canonical ISGs (49, 50). Of note, Type-III IFN appears to activate a network of genes most similar to that of Type-I IFNs (51). Future investigations are needed to address the effects of Type-III IFN on uterine macrophages.

While both Type-I and Type-II IFNs are capable of directing macrophages to upregulate CD122 in vitro, it remains to be determined which IFNs do so in vivo. It is possible that several different interferons at several different timepoints in pregnancy act on macrophages, as observed developmental kinetics of CD122+Macs mirrored that of total IFN activity in the uterus and placenta (30). Mouse and human uterine glandular epithelial cells are an established reservoir of Type-I IFN (52, 53). For example, the Type-I IFNε is produced by uterine epithelium under direction of estrogen (53). While a population of CD122+Macs exists in the non-pregnant uterus of IFNAR KO mice under direction of M-CSF and a non-Type-I IFN and/or additional signal, IFNε may still contribute to development of CD122+Macs in the normal non-pregnant uterus.

Regulation of IFNγ during murine pregnancy has been thoroughly described (54). Levels progressively rise to a peak at mid-gestation, after which levels halve but plateau to near-term. The cells upon which we performed gene expression profiling were obtained on E7.5, when levels of uterine IFNγ are known to be relatively low. Consistent with those data, gene expression and gene ontology analyses of CD122+Macs appeared most consistent with a Type-I or -III IFN signature. The transcriptome of CD122+Macs was dominated by ISGs, including *Mx1*, that are induced preferentially in response to IFN-I and -III (49, 55). Further, CD122+Macs are largely MHCII_low_ in vivo. Type-II IFN has been shown to enhance expression of MHCII in antigen presenting cells, while IFN-I has been shown to oppose IFN-II-mediated upregulation of MHCII (34). We also showed that treatment of BMDMs with IFNγ in vitro led to upregulation of CD122 and MHCII, while treatment of BMDMs with IFNα led to upregulation of CD122 but not MHCII. The IFN system is complex and redundant, particularly the Type-I and Type-III families of IFNs, each composed of several members. This makes isolating individual actors in vivo difficult. Future work will be directed at dissecting this complex system to determine the in vivo IFN requirements for generation of CD122+Macs during preganacy.

We observed that only macrophages, not other uterine myeloid cells, express CD122. Further, only CD122+Macs, not CD122+ NK cells, experienced a boost in expression of CD122 on a per-cell basis that correlated with increased IFN activity during gestation. One interpretation is that only macrophages are in close enough proximity to the local source of IFN to express CD122 under direction of IFNs. An alternative explanation is that upregulation of CD122 in response to IFN is a mechanism unique to monocytes and macrophages. In support of this latter hypothesis, we provided evidence in vitro that M-CSF-derived macrophages, not GM-CSF-derived macrophages, responded to IFN by upregulating CD122. These data point to unique regulation of *Il2rb* in macrophages, allowing it to be expressed rapidly upon signaling by IFNs. Relatively little is known about transcriptional regulation of *Il2rb* (56, 57), and further investigations into why macrophages uniquely respond to IFN by upregulating CD122 are needed. This is especially true in light of the fact that CD122+Macs express neither Eomes nor T-bet, which have been previously shown to drive expression of *Il2rb* in killer lymphocytes (58). In summary, our data support the notion that CD122 is a novel, macrophage-specific, IFN-stimulated gene.

Our data show that CD122+Macs can derive from BM monocytes, but the precise precursors of CD122+Macs have yet to be determined. We found evidence of CD122+F4/80_int_ cells in the non-pregnant uterus that likely represent CD122+ monocytes. These cells may represent an intermediate through which CD122+Macs transit during gestation. It has long been appreciated that M-CSF is critical for pregnancy and is positively regulated by sex hormones (43, 59, 60). M-CSF concentrations are at or below the limits of detection in the uterus of non-pregnant mice but increase dramatically in the myometrium during gestation, reaching peak concentration at term (26, 59, 60). Indeed, we showed that large, F4/80_bright_ macrophages were absent from the non-pregnant uterus. The appearance of these cells in the pregnant uterus suggests they may derive from uterine monocytes exposed to M-CSF. In contrast to the myometrium, concentration of M-CSF is modest and constant in the decidua and placenta as gestation progresses (59). Reinforcing these data, additional work showed maintenance of myeloid cells in the growing myometrium and decline of myeloid populations in the decidua as gestation progresses (26, 27). These data are consistent with our observations that adoptively transferred monocytes converted into F4/80_bright_CD122+Macs in the myometrium but not the decidua or placenta late in gestation (data not shown). Altogether, these data are consistent with a model in which BM monocytes are directed to develop into F4/80_bright_ CD122+Macs by M-CSF and IFNs.

Finally, we found that CD122+Macs exhibit a biochemical and functional response to stimulation with IL-15. Whether CD122+Macs respond to IL-15 in the same manner as classical IL-15-responsive killer lymphocytes, however, remains to be determined. It has long been appreciated that IL-15 enhances cytotoxicity of NK cells (61), and we found it intruiging that CD122+Macs expressed a number of cytolytic transcripts. IL-15 may be responsible for the expression of granzymes and perforin observed in CD122+Macs, as IL-15 has been implicated in driving expression of cytolytic mediators in human plasma cells (62), another non-traditional responder to IL-15.

We also have yet to understand the specific stimuli responsible for activating uterine and placental macrophages in vivo during pregnancy, but it is feasible that CD122+Macs respond to IL-15 by enhancing production of proinflammatory cytokines that may favor or threaten a healthy pregnancy. For instance, decidual CD122+Macs were enriched in *Il18* transcript (encoding IL-18), which is known to support production of IFNγ at the maternal-fetal interface. Establishing this link may have major clinical relevance. Mice deficient in IFNγ signaling fail to remodel uterine spiral arteries into low-resistance, high-capacitance vessels (12). In the mouse, this has been attributed to defective production of IFNγ by NK cells. Further, IL-12 and IL-18 appear to play a key role in production of IFNγ at the maternal fetal interface, as mice deficient in either or both of these cytokines exhibit similarly abnormal uterine artery remodeling as does the IFNγ KO mouse (63). Thus, these mice develop the correlate of human preeclampsia (12, 64). At the same time, hyper-stimulation of CD122+Macs with an over-abundance of IL-15 may contribute to similar adverse outcomes of pregnancy, as seen in a mouse model of spontaneous preeclampsia characterized by an abundance of IL-15 and a relative paucity of NK cells (65). Investigations into the IL-15-dependent and IL-15-independent functions of CD122+Macs in pregnancy are ongoing and may shed light on new therapies for preeclampsia.

Lending validation to our findings in the mouse, we discovered that the human uterus contains a population of CD122+Macs. We observed that these cells phenocopied mouse CD122+Macs, with expression of CD122 and modest levels of NK cell markers, such as CD56, Perforin, and Granzyme. Human CD122+Macs were not as abundant as mouse CD122+Macs early in gestation, but we have yet to fully explore how this population changes over gestation in humans. We also have yet to formally determine whether these human womb-associated, killer-like macrophages develop, signal, and function like their murine counterparts. A prior report supports the notion that IFNs may be able to drive expression of CD122 in human macrophages, as cultured human blood monocytes can express CD122 after culture in high-dose IFNγ (66). Our findings greatly extend and provide critical context for these prior data. How IFNs signal to human monocytes and macrophages is a key area of ongoing investigation with clear implications for human pregnancy. In summary, we have revealed that IFNs act on uterine macrophages to generate an entirely new and unexpected IL-15-responsive cell type at the maternal-fetal interface, the CD122+ macrophage. Given the importance of IL-15 in pregnancy, modulation of CD122+Macs may represent a novel therapeutic target to support pregnancies threatened by fetal growth restriction, fetal loss, and preeclampsia.

## ACKNOWLEDGMENTS

None.

The authors have declared that no conflict of interest exists.

