## Supplemental for "Interferons drive development of novel interleukin-15-responsive macrophages"

### SUPPLEMENTAL FIGURE 1

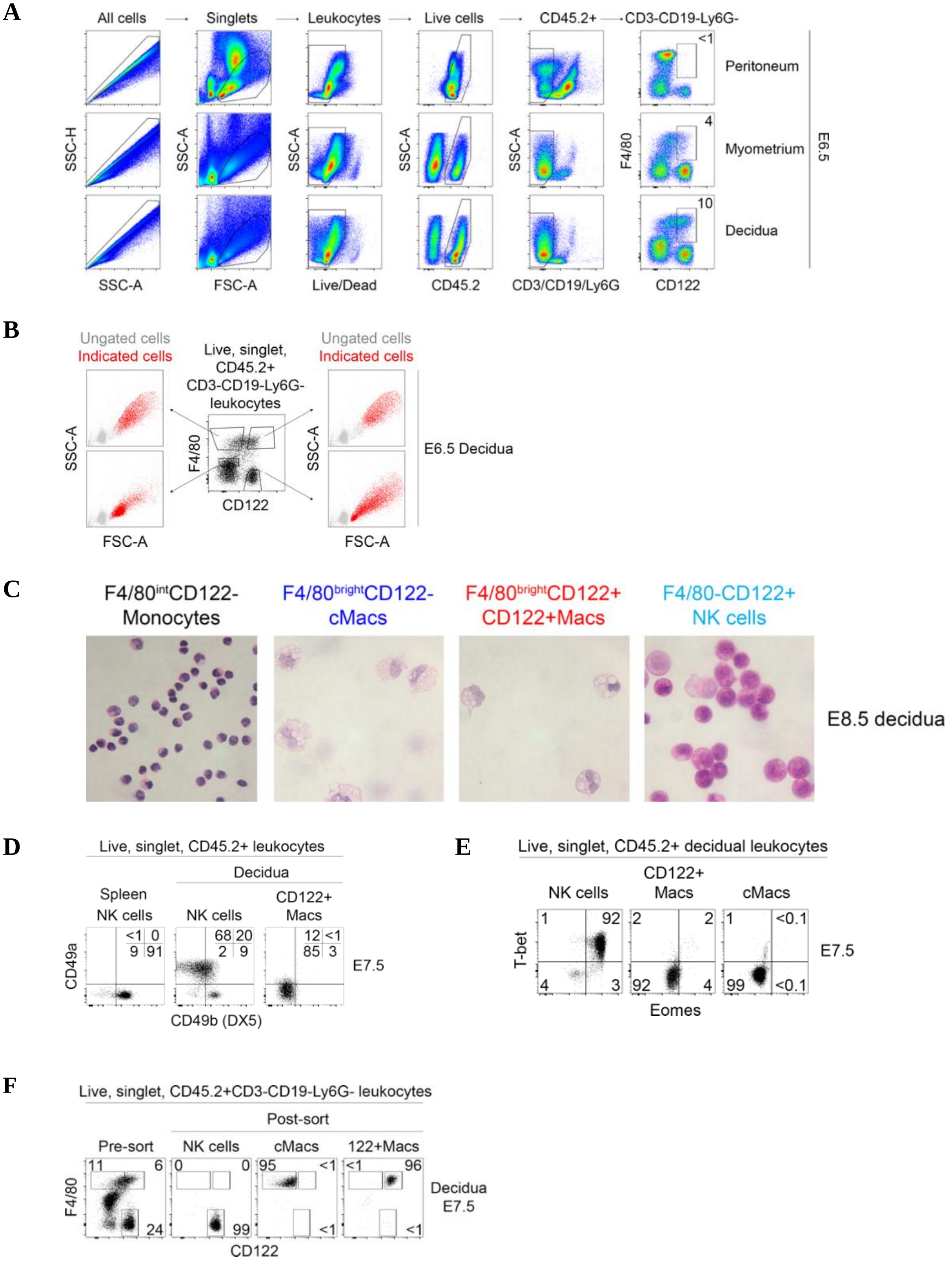

**Supplemental Figure 1. CD122+Macs are not NK cells, dendritic cells, or neutrophils.** (A) Shown is the raw gating scheme used to identify murine CD122+Macs. Data are representative of at least 20 independent experiments. (B) Like cMacs (F4/80$_{bright}$CD122-), CD122+Macs (F4/80$_{bright}$CD122+) are among the largest cells in the decidua at E6.5 by flow cytometric analysis. (C) From left, decidual CD122-F4/80$_{int}$ cells at E8.5 are morphologically monocyte-like, all F4/80$_{bright}$ cells are vacuolated and macrophage-like, and CD122+F4/80- cells are more finely granular and NK-like. Indicated cells were FACS-sorted and stained with hematoxylin and eosin. Data are representative of 2 independent experiments. (D) Conventional splenic NK cells are CD49b/DX5+, decidual NK cells are largely CD49a+, and CD122+Macs express neither CD49a nor CD49b. Data are representative of 2-3 independent experiments with 2-3 mice per experiment. (E) Similarly, decidual NK cells express high levels of both T-bet and Eomes, while neither CD122+Macs nor cMacs express T-bet or Eomes. Data are representative of 2-3 independent experiments with 2-3 mice per experiment. (F) Shown is the sorting strategy to assess transcriptomes of purified decidual CD122+Macs, cMacs, and NK cells. Cells to be profiled by microarray were FACS-sorted to at least 95% purity from 3 independent groups of pooled E7.5 deciduae, with each group consisting of 4-5 mice.

| Gene symbol | Gene name | Absolute fold change (CD122+Macs/NK) | Adjusted *p* value |
|---|---|---|---|
| C3ar1 | complement component 3a receptor 1 | 302.56 | 8.29E-13 |
| Pf4 | platelet factor 4 | 249.69 | 1.74E-12 |
| C1qa | complement component 1, q subcomponent, alpha polypeptide | 211.69 | 1.23E-10 |
| C1qb | complement component 1, q subcomponent, beta polypeptide | 200.13 | 3.16E-10 |
| Ccl3 | chemokine (C-C motif) ligand 3 | 173.44 | 1.14E-10 |
| Ms4a7 | membrane-spanning 4-domains, subfamily A, member 7 | 171.44 | 1.62E-12 |
| C1qc | complement component 1, q subcomponent, C chain | 167.21 | 2.02E-12 |
| Mrc1 | mannose receptor, C type 1 | 159.25 | 3.09E-11 |
| Ly86 | lymphocyte antigen 86 | 158.62 | 3.40E-12 |
| Fcgr4 | Fc receptor, IgG, low affinity IV | 157.70 | 1.06E-11 |
| Pld4 | phospholipase D family, member 4 | 151.62 | 1.13E-11 |
| Lyz2 | lysozyme 2 | 147.95 | 1.51E-10 |
| Cbr2 | carbonyl reductase 2 | 144.50 | 4.39E-12 |
| Ms4a6d | membrane-spanning 4-domains, subfamily A, member 6D | 143.47 | 5.60E-12 |
| Ccl8 | chemokine (C-C motif) ligand 8 | 141.11 | 1.27E-10 |
| Tlr7 | toll-like receptor 7 | 112.90 | 1.16E-11 |
| Cybb | cytochrome b-245, beta polypeptide | 108.59 | 3.11E-11 |
| Ctsh | cathepsin H | 104.16 | 3.40E-12 |
| Ifi27l2a | interferon, alpha-inducible protein 27 like 2A | 102.68 | 1.06E-11 |
| Ms4a6c | membrane-spanning 4-domains, subfamily A, member 6C | 101.60 | 5.40E-12 |
| Sirpa | signal-regulatory protein alpha | 100.59 | 1.01E-10 |
| Fcrls | Fc receptor-like S, scavenger receptor | 100.10 | 3.40E-12 |
| Ms4a14 | membrane-spanning 4-domains, subfamily A, member 14 | 100.03 | 5.40E-12 |
| Ccl2 | chemokine (C-C motif) ligand 2 | 99.48 | 2.02E-12 |
| Mpeg1 | macrophage expressed gene 1 | 98.61 | 1.25E-10 |
| Tlr1 | toll-like receptor 1 | 94.76 | 4.31E-12 |
| Ifi207 | interferon activated gene 207 | 91.95 | 4.12E-10 |
| Lgmn | legumain | 86.64 | 1.33E-09 |
| Fcgr1 | Fc receptor, IgG, high affinity I | 86.31 | 3.40E-12 |
| Ccl6 | chemokine (C-C motif) ligand 6 | 76.14 | 2.05E-11 |
| Cxcl16 | chemokine (C-X-C motif) ligand 16 | 76.12 | 3.42E-11 |
| Cxcl2 | chemokine (C-X-C motif) ligand 2 | 74.25 | 5.60E-12 |
| Tlr13 | toll-like receptor 13 | 73.81 | 5.60E-12 |
| Blnk | B cell linker | 72.14 | 5.60E-12 |
| Adgre1 | adhesion G protein-coupled receptor E1 | 68.94 | 1.89E-11 |
| Dab2 | disabled 2, mitogen-responsive phosphoprotein | 64.81 | 4.24E-11 |
| P2ry6 | pyrimidinergic receptor P2Y, G-protein coupled, 6 | 63.89 | 1.27E-10 |
| P2rx4 | purinergic receptor P2X, ligand-gated ion channel 4 | 63.22 | 1.13E-11 |
| Gm6377 | predicted gene 6377 | 62.86 | 2.12E-11 |
| C5ar1 | complement component 5a receptor 1 | 61.92 | 6.28E-11 |
| Alox5ap | arachidonate 5-lipoxygenase activating protein | 61.47 | 8.27E-11 |
| Clec4n | C-type lectin domain family 4, member n | 56.67 | 2.00E-10 |
| Lst1 | leukocyte specific transcript 1 | 56.29 | 2.33E-10 |
| Apoe | apolipoprotein E | 56.07 | 3.11E-11 |
| Aif1 | allograft inflammatory factor 1 | 55.55 | 4.21E-11 |
| Tlr8 | toll-like receptor 8 | 50.64 | 1.40E-11 |
| Wfdc17 | WAP four-disulfide core domain 17 | 50.59 | 9.21E-10 |
| Ccl7 | chemokine (C-C motif) ligand 7 | 50.54 | 7.01E-11 |
| Clec4a1 | C-type lectin domain family 4, member a1 | 50.24 | 2.02E-11 |
| Il1b | interleukin 1 beta | 49.90 | 8.89E-11 |
| Cd74 | CD74 antigen (invariant polypeptide of major histocompatibility complex, class II antigen-associated) | 49.50 | 5.81E-11 |
| Cd300c2 | CD300C molecule 2 | 47.99 | 2.34E-10 |
| Cd180 | CD180 antigen | 47.75 | 1.29E-10 |
| Cx3cr1 | chemokine (C-X3-C motif) receptor 1 | 47.47 | 4.96E-10 |

| Gene symbol | Gene name | Absolute fold change (CD122+Macs/NK) | Adjusted *p* value |
|---|---|---|---|
| Csf1r | colony stimulating factor 1 receptor | 47.44 | 5.43E-10 |
| Plbd1 | phospholipase B domain containing 1 | 46.96 | 1.13E-11 |
| Cd86 | CD86 antigen | 46.65 | 1.16E-11 |
| Ms4a4c | membrane-spanning 4-domains, subfamily A, member 4C | 46.59 | 4.47E-11 |
| Tlr2 | toll-like receptor 2 | 46.54 | 2.48E-11 |
| Fcgr2b | Fc receptor, IgG, low affinity IIb | 46.51 | 9.70E-10 |
| Ms4a4a | membrane-spanning 4-domains, subfamily A, member 4A | 45.83 | 5.24E-09 |
| Il18 | interleukin 18 | 45.27 | 3.15E-11 |
| Ifit2 | interferon-induced protein with tetratricopeptide repeats 2 | 45.07 | 7.01E-11 |
| Slc15a3 | solute carrier family 15, member 3 | 44.62 | 1.28E-09 |
| Cd33 | CD33 antigen | 44.02 | 3.26E-11 |
| Apobec1 | apolipoprotein B mRNA editing enzyme, catalytic polypeptide 1 | 44.00 | 4.47E-11 |
| Ifngr2 | interferon gamma receptor 2 | 43.35 | 2.40E-10 |
| Ifit3 | interferon-induced protein with tetratricopeptide repeats 3 | 42.46 | 2.02E-11 |
| H2-Aa | histocompatibility 2, class II antigen A, alpha | 41.96 | 9.72E-12 |
| Gpr137b-ps | G protein-coupled receptor 137B, pseudogene | 41.90 | 6.67E-12 |
| Hpgds | hematopoietic prostaglandin D synthase | 41.69 | 1.16E-11 |
| Spi1 | spleen focus forming virus (SFFV) proviral integration oncogene | 40.74 | 3.42E-11 |
| Egr2 | early growth response 2 | 37.33 | 1.45E-10 |
| Oas2 | 2'-5' oligoadenylate synthetase 2 | 36.75 | 6.52E-10 |
| G530011O06Rik | RIKEN cDNA G530011O06 gene | 36.73 | 1.86E-10 |
| Igsf6 | immunoglobulin superfamily, member 6 | 36.00 | 4.45E-11 |
| Tmem106a | transmembrane protein 106A | 35.26 | 2.33E-10 |
| Slfn1 | schlafen 1 | 35.07 | 8.52E-10 |
| H2-Ab1 | histocompatibility 2, class II antigen A, beta 1 | 34.34 | 1.05E-09 |
| Hck | hemopoietic cell kinase | 33.67 | 8.21E-11 |
| Lpcat2 | lysophosphatidylcholine acyltransferase 2 | 33.66 | 1.71E-11 |
| Clec4a2 | C-type lectin domain family 4, member a2 | 33.52 | 2.05E-09 |
| Clec7a | C-type lectin domain family 7, member a | 33.11 | 2.17E-10 |
| Cd68 | CD68 antigen | 32.74 | 1.73E-10 |
| Folr2 | folate receptor 2 (fetal) | 32.54 | 6.48E-11 |
| Rab7b | RAB7B, member RAS oncogene family | 32.50 | 1.20E-10 |
| Ccl12 | chemokine (C-C motif) ligand 12 | 32.44 | 8.89E-11 |
| Slc11a1 | solute carrier family 11 (proton-coupled divalent metal ion transporters), member 1 | 31.70 | 1.89E-09 |
| Milr1 | mast cell immunoglobulin like receptor 1 | 31.41 | 1.79E-10 |
| Clec12a | C-type lectin domain family 12, member a | 31.13 | 1.98E-10 |
| Gm2a | GM2 ganglioside activator protein | 30.64 | 4.47E-11 |
| Tgfbi | transforming growth factor, beta induced | 30.27 | 1.71E-11 |
| Sdc3 | syndecan 3 | 30.23 | 1.51E-09 |
| Cd36 | CD36 molecule | 30.20 | 3.14E-10 |
| Rab3il1 | RAB3A interacting protein (rabin3)-like 1 | 30.03 | 3.14E-09 |
| Siglec1 | sialic acid binding Ig-like lectin 1, sialoadhesin | 29.87 | 1.30E-09 |
| Cd300ld | CD300 molecule like family member d | 29.46 | 3.42E-11 |
| Stab1 | stabilin 1 | 29.22 | 6.66E-10 |
| Rgs10 | regulator of G-protein signalling 10 | 28.95 | 1.60E-11 |
| Thbs1 | thrombospondin 1 | 28.79 | 1.00E-09 |
| Itih5 | inter-alpha (globulin) inhibitor H5 | 0.13 | 2.68E-05 |
| Styk1 | serine/threonine/tyrosine kinase 1 | 0.13 | 7.27E-07 |
| Ppm1j | protein phosphatase 1J | 0.13 | 3.53E-08 |
| Gimap9 | GTPase, IMAP family member 9 | 0.13 | 2.85E-08 |
| Itk | IL2 inducible T cell kinase | 0.13 | 1.09E-08 |
| Gem | GTP binding protein (gene overexpressed in skeletal muscle) | 0.13 | 1.55E-06 |
| Gipc2 | GIPC PDZ domain containing family, member 2 | 0.12 | 1.51E-08 |
| Bhlhe40 | basic helix-loop-helix family, member e40 | 0.12 | 4.66E-08 |

| Gene symbol | Gene name | Absolute fold change (CD122+Macs/NK) | Adjusted *p* value |
|---|---|---|---|
| Epdr1 | ependymin related protein 1 (zebrafish) | 0.12 | 6.85E-08 |
| Rhof | ras homolog family member F (in filopodia) | 0.12 | 6.82E-07 |
| Sept1 | septin 1 | 0.12 | 9.70E-08 |
| Tnfrsf18 | tumor necrosis factor receptor superfamily, member 18 | 0.12 | 4.83E-06 |
| Klra3 | killer cell lectin-like receptor, subfamily A, member 3 | 0.11 | 6.41E-09 |
| Stk39 | serine/threonine kinase 39 | 0.11 | 6.21E-09 |
| F2r | coagulation factor II (thrombin) receptor | 0.11 | 3.58E-09 |
| Stk26 | serine/threonine kinase 26 | 0.11 | 6.02E-09 |
| Rnf125 | ring finger protein 125 | 0.11 | 2.27E-08 |
| D16Ertd472e | DNA segment, Chr 16, ERATO Doi 472, expressed | 0.11 | 1.33E-09 |
| Gm24463 | predicted gene, 24463 | 0.11 | 3.46E-09 |
| Car5b | carbonic anhydrase 5b, mitochondrial | 0.11 | 8.50E-09 |
| Zap70 | zeta-chain (TCR) associated protein kinase | 0.11 | 9.55E-08 |
| Prkcq | protein kinase C, theta | 0.10 | 1.97E-09 |
| Ablim1 | actin-binding LIM protein 1 | 0.10 | 1.10E-08 |
| Ppp3cc | protein phosphatase 3, catalytic subunit, gamma isoform | 0.10 | 6.53E-10 |
| Fam174b | family with sequence similarity 174, member B | 0.10 | 1.97E-07 |
| Gimap6 | GTPase, IMAP family member 6 | 0.10 | 6.07E-09 |
| Dsc2 | desmocollin 2 | 0.10 | 1.25E-06 |
| Satb1 | special AT-rich sequence binding protein 1 | 0.10 | 3.46E-09 |
| Sidt1 | SID1 transmembrane family, member 1 | 0.10 | 2.37E-08 |
| Pecam1 | platelet/endothelial cell adhesion molecule 1 | 0.09 | 2.70E-09 |
| Gpr141 | G protein-coupled receptor 141 | 0.09 | 4.61E-09 |
| Gm40275 | predicted gene, 40275 | 0.09 | 4.43E-09 |
| Klrk1 | killer cell lectin-like receptor subfamily K, member 1 | 0.09 | 8.97E-10 |
| Avil | advillin | 0.09 | 9.29E-10 |
| Khdc1a | KH domain containing 1A | 0.08 | 3.74E-10 |
| Cd3g | CD3 antigen, gamma polypeptide | 0.08 | 1.95E-09 |
| Pglyrp1 | peptidoglycan recognition protein 1 | 0.08 | 6.18E-09 |
| Ifitm1 | interferon induced transmembrane protein 1 | 0.08 | 2.07E-09 |
| Serpinb9 | serine (or cysteine) peptidase inhibitor, clade B, member 9 | 0.08 | 1.71E-09 |
| I730030J21Rik | RIKEN cDNA I730030J21 gene | 0.08 | 1.37E-08 |
| Myb | myeloblastosis oncogene | 0.08 | 1.53E-09 |
| Gimap7 | GTPase, IMAP family member 7 | 0.07 | 3.22E-09 |
| Sult2b1 | sulfotransferase family, cytosolic, 2B, member 1 | 0.07 | 1.91E-08 |
| Tbx21 | T-box 21 | 0.07 | 6.56E-10 |
| Scin | scinderin | 0.07 | 2.03E-07 |
| Klra21 | killer cell lectin-like receptor subfamily A, member 21 | 0.07 | 2.59E-09 |
| Sytl3 | synaptotagmin-like 3 | 0.07 | 7.50E-10 |
| Atp1b1 | ATPase, Na+/K+ transporting, beta 1 polypeptide | 0.06 | 4.13E-06 |
| Sla2 | Src-like-adaptor 2 | 0.06 | 5.89E-07 |
| Samd3 | sterile alpha motif domain containing 3 | 0.06 | 8.21E-11 |
| Gzmd | granzyme D | 0.06 | 1.37E-09 |
| P2ry10 | purinergic receptor P2Y, G-protein coupled 10 | 0.06 | 3.21E-08 |
| Ctsg | cathepsin G | 0.06 | 1.75E-09 |
| Ptprcap | protein tyrosine phosphatase, receptor type, C polypeptide-associated protein | 0.06 | 4.13E-09 |
| Zbtb32 | zinc finger and BTB domain containing 32 | 0.06 | 1.97E-07 |
| Ptpn3 | protein tyrosine phosphatase, non-receptor type 3 | 0.06 | 1.85E-09 |
| Myo10 | myosin X | 0.06 | 1.89E-09 |
| Il2ra | interleukin 2 receptor, alpha chain | 0.05 | 1.48E-08 |
| Fasl | Fas ligand (TNF superfamily, member 6) | 0.05 | 4.44E-11 |
| Gzmg | granzyme G | 0.05 | 9.41E-09 |
| Icos | inducible T cell co-stimulator | 0.05 | 6.69E-10 |
| Lax1 | lymphocyte transmembrane adaptor 1 | 0.05 | 3.49E-10 |

| Gene symbol | Gene name | Absolute fold change (CD122+Macs/NK) | Adjusted *p* value |
|---|---|---|---|
| Clnk | cytokine-dependent hematopoietic cell linker | 0.05 | 3.53E-11 |
| Gimap4 | GTPase, IMAP family member 4 | 0.05 | 1.99E-10 |
| Srgap3 | SLIT-ROBO Rho GTPase activating protein 3 | 0.05 | 1.75E-10 |
| Zdhhc15 | zinc finger, DHHC domain containing 15 | 0.05 | 5.24E-11 |
| Gzmb | granzyme B | 0.05 | 2.80E-09 |
| Cst7 | cystatin F (leukocystatin) | 0.05 | 9.70E-10 |
| Itga1 | integrin alpha 1 | 0.04 | 1.30E-09 |
| Gzmc | granzyme C | 0.04 | 6.56E-10 |
| E330009J07Rik | RIKEN cDNA E330009J07 gene | 0.04 | 1.60E-09 |
| Sh2d1b2 | SH2 domain containing 1B2 | 0.04 | 8.38E-11 |
| Ctsw | cathepsin W | 0.04 | 4.81E-11 |
| Klri2 | killer cell lectin-like receptor family I member 2 | 0.04 | 3.11E-11 |
| Prf1 | perforin 1 (pore forming protein) | 0.04 | 3.35E-09 |
| Klrb1b | killer cell lectin-like receptor subfamily B member 1B | 0.03 | 5.68E-11 |
| Txk | TXK tyrosine kinase | 0.03 | 1.00E-09 |
| Car2 | carbonic anhydrase 2 | 0.03 | 4.39E-11 |
| Eomes | eomesodermin | 0.03 | 6.07E-11 |
| Sytl2 | synaptotagmin-like 2 | 0.03 | 1.65E-10 |
| Klrb1c | killer cell lectin-like receptor subfamily B member 1C | 0.03 | 5.16E-10 |
| Klra5 | killer cell lectin-like receptor, subfamily A, member 5 | 0.03 | 1.50E-10 |
| Gzma | granzyme A | 0.03 | 1.51E-10 |
| Cd7 | CD7 antigen | 0.03 | 1.59E-10 |
| Sh2d1a | SH2 domain containing 1A | 0.03 | 8.89E-11 |
| Sh2d2a | SH2 domain containing 2A | 0.03 | 5.97E-09 |
| Thy1 | thymus cell antigen 1, theta | 0.02 | 7.62E-11 |
| Gpr87 | G protein-coupled receptor 87 | 0.02 | 2.48E-11 |
| Tnfrsf9 | tumor necrosis factor receptor superfamily, member 9 | 0.02 | 2.02E-11 |
| Cd96 | CD96 antigen | 0.02 | 6.60E-12 |
| Tmsb15a | thymosin beta 15a | 0.02 | 1.40E-11 |
| Mcpt8 | mast cell protease 8 | 0.02 | 1.37E-11 |
| Ctla4 | cytotoxic T-lymphocyte-associated protein 4 | 0.02 | 1.45E-10 |
| Il2rb | interleukin 2 receptor, beta chain | 0.02 | 6.56E-09 |
| Xcl1 | chemokine (C motif) ligand 1 | 0.02 | 3.09E-11 |
| Klrg1 | killer cell lectin-like receptor subfamily G, member 1 | 0.02 | 5.43E-10 |
| Klre1 | killer cell lectin-like receptor family E member 1 | 0.01 | 7.23E-12 |
| Klrd1 | killer cell lectin-like receptor, subfamily D, member 1 | 0.01 | 5.13E-12 |
| Nkg7 | natural killer cell group 7 sequence | 0.01 | 1.71E-11 |
| Ncr1 | natural cytotoxicity triggering receptor 1 | 0.01 | 4.39E-12 |

**Supplemental Table 1. Macrophage-associated transcripts are abundant in decidual CD122+Macs, while killer lymphocyte transcripts are abundant in decidual NK cells.** Changes in gene expression by microarray between sort-purified CD122+Macs and NK cells. Only the top 200 differentially expressed genes by fold change (100 enriched in CD122+Macs and 100 enriched in NK cells) with an adjusted *p* value $<0.05$ are shown. Absolute fold change of each individual gene refers to level in CD122+Macs relative to level in NK cells. Of note, *Il2rb* (encoding CD122) was highly enriched in NK cells despite modest to no difference in detection of surface CD122 by flow cytometry.

| Gene symbol | Gene name | Absolute fold change (CD122+Macs/cMacs) | Adjusted *p* value |
|---|---|---|---|
| Ccl8 | chemokine (C-C motif) ligand 8 | 4.16 | 5.75E-04 |
| Oas3 | 2'-5' oligoadenylate synthetase 3 | 3.90 | 7.57E-04 |
| Cfb | complement factor B | 3.47 | 1.04E-04 |
| Ifi205 | interferon activated gene 205 | 3.35 | 3.69E-04 |
| Ifit2 | interferon-induced protein with tetratricopeptide repeats 2 | 3.22 | 3.06E-04 |
| Mmp27 | matrix metallopeptidase 27 | 3.05 | 5.55E-03 |
| Zbp1 | Z-DNA binding protein 1 | 2.92 | 1.04E-04 |
| Irf7 | interferon regulatory factor 7 | 2.90 | 2.77E-02 |
| Gzmc | granzyme C | 2.86 | 7.57E-04 |
| Oasl1 | 2'-5' oligoadenylate synthetase-like 1 | 2.85 | 4.94E-04 |
| Gzmb | granzyme B | 2.84 | 2.28E-03 |
| Ifit3b | interferon-induced protein with tetratricopeptide repeats 3B | 2.82 | 1.30E-04 |
| Ifi44 | interferon-induced protein 44 | 2.73 | 9.05E-04 |
| Fabp5 | fatty acid binding protein 5, epidermal | 2.72 | 4.94E-04 |
| Fabp3 | fatty acid binding protein 3, muscle and heart | 2.68 | 4.93E-04 |
| Slfn1 | schlafen 1 | 2.67 | 3.17E-03 |
| Spp1 | secreted phosphoprotein 1 | 2.66 | 5.52E-04 |
| Ifi208 | interferon activated gene 208 | 2.59 | 2.65E-03 |
| Gzmd | granzyme D | 2.51 | 1.87E-03 |
| Cd300lf | CD300 molecule like family member F | 2.48 | 5.63E-04 |
| Phf11a | PHD finger protein 11A | 2.41 | 9.63E-03 |
| Tnfsf10 | tumor necrosis factor (ligand) superfamily, member 10 | 2.40 | 2.47E-03 |
| Rsad2 | radical S-adenosyl methionine domain containing 2 | 2.39 | 3.72E-03 |
| Hpse | heparanase | 2.36 | 4.51E-04 |
| Ifit3 | interferon-induced protein with tetratricopeptide repeats 3 | 2.33 | 5.75E-04 |
| Atp6v0d2 | ATPase, H+ transporting, lysosomal V0 subunit D2 | 2.29 | 2.41E-03 |
| Cmpk2 | cytidine monophosphate (UMP-CMP) kinase 2, mitochondrial | 2.24 | 2.28E-03 |
| Ms4a4c | membrane-spanning 4-domains, subfamily A, member 4C | 2.23 | 2.41E-03 |
| Ifit1 | interferon-induced protein with tetratricopeptide repeats 1 | 2.23 | 4.69E-03 |
| Isg20 | interferon-stimulated protein | 2.22 | 1.02E-03 |
| Nt5c3 | 5'-nucleotidase, cytosolic III | 2.21 | 6.75E-04 |
| Gpnmb | glycoprotein (transmembrane) nmb | 2.17 | 2.64E-03 |
| Gbp8 | guanylate-binding protein 8 | 2.16 | 3.83E-03 |
| Cd200r4 | CD200 receptor 4 | 2.16 | 1.65E-02 |
| Pde7b | phosphodiesterase 7B | 2.15 | 4.08E-03 |
| Slfn4 | schlafen 4 | 2.14 | 3.79E-03 |
| Gbp7 | guanylate binding protein 7 | 2.14 | 4.18E-02 |
| Usp18 | ubiquitin specific peptidase 18 | 2.13 | 2.25E-03 |
| Siglec1 | sialic acid binding Ig-like lectin 1, sialoadhesin | 2.13 | 1.66E-02 |
| Gdf15 | growth differentiation factor 15 | 2.12 | 2.41E-03 |
| Dhx58 | DEXH (Asp-Glu-X-His) box polypeptide 58 | 2.12 | 6.95E-03 |
| Irgm1 | immunity-related GTPase family M member 1 | 2.09 | 1.81E-02 |
| Phf11d | PHD finger protein 11D | 2.07 | 9.42E-03 |
| Ifi209 | interferon activated gene 209 | 2.06 | 3.72E-03 |
| Gm5431 | predicted gene 5431 | 2.05 | 3.51E-02 |
| Gzmg | granzyme G | 2.04 | 3.42E-02 |
| Ifi206 | interferon activated gene 206 | 2.04 | 3.29E-03 |
| BC147527 | cDNA sequence BC147527 | 2.02 | 6.34E-03 |
| Gm12250 | predicted gene 12250 | 2.00 | 1.73E-03 |
| Cbr2 | carbonyl reductase 2 | 1.98 | 3.20E-03 |
| Gbp4 | guanylate binding protein 4 | 1.98 | 6.17E-03 |
| Stat1 | signal transducer and activator of transcription 1 | 1.97 | 4.40E-02 |
| Gm1966 | predicted gene 1966 | 1.96 | 2.07E-02 |
| Phf11c | PHD finger protein 11C | 1.95 | 3.72E-03 |
| Gbp9 | guanylate-binding protein 9 | 1.95 | 2.41E-03 |
| Slc9a7 | solute carrier family 9 (sodium/hydrogen exchanger), member 7 | 1.95 | 7.76E-03 |
| Nkg7 | natural killer cell group 7 sequence | 1.92 | 7.07E-03 |
| Clec10a | C-type lectin domain family 10, member A | 1.92 | 4.95E-02 |
| Treml2 | triggering receptor expressed on myeloid cells-like 2 | 1.90 | 5.67E-03 |
| Ddx58 | DEAD (Asp-Glu-Ala-Asp) box polypeptide 58 | 1.89 | 2.43E-02 |
| Ddx60 | DEAD (Asp-Glu-Ala-Asp) box polypeptide 60 | 1.88 | 2.39E-02 |
| Slfn2 | schlafen 2 | 1.88 | 2.87E-02 |
| Msrb1 | methionine sulfoxide reductase B1 | 1.88 | 1.84E-02 |
| Xdh | xanthine dehydrogenase | 1.86 | 4.18E-02 |

| Gene symbol | Gene name | Absolute fold change (CD122+Macs/cMacs) | Adjusted *p* value |
|---|---|---|---|
| Npl | N-acetylneuraminate pyruvate lyase | 1.86 | 1.14E-02 |
| Lrp12 | low density lipoprotein-related protein 12 | 1.85 | 2.82E-03 |
| Ube2l6 | ubiquitin-conjugating enzyme E2L 6 | 1.85 | 1.13E-02 |
| Gas2l3 | growth arrest-specific 2 like 3 | 1.81 | 3.85E-02 |
| Mx1 | MX dynamin-like GTPase 1 | 1.80 | 4.08E-03 |
| G530011O06Rik | RIKEN cDNA G530011O06 gene | 1.80 | 2.88E-02 |
| Parp11 | poly (ADP-ribose) polymerase family, member 11 | 1.79 | 8.69E-03 |
| Agtrap | angiotensin II, type I receptor-associated protein | 1.79 | 2.60E-02 |
| Papss2 | 3'-phosphoadenosine 5'-phosphosulfate synthase 2 | 1.78 | 1.43E-02 |
| Gbp5 | guanylate binding protein 5 | 1.78 | 5.44E-03 |
| Tmem140 | transmembrane protein 140 | 1.77 | 1.00E-02 |
| Stat2 | signal transducer and activator of transcription 2 | 1.77 | 1.35E-02 |
| Dtx3l | deltex 3-like, E3 ubiquitin ligase | 1.77 | 4.14E-02 |
| Nxpe5 | neurexophilin and PC-esterase domain family, member 5 | 1.76 | 1.95E-02 |
| Tgm2 | transglutaminase 2, C polypeptide | 1.74 | 1.55E-02 |
| Tmsb15a | thymosin beta 15a | 1.73 | 7.68E-03 |
| Fgl2 | fibrinogen-like protein 2 | 1.73 | 5.67E-03 |
| H2-T24 | histocompatibility 2, T region locus 24 | 1.72 | 2.66E-02 |
| Clcn7 | chloride channel, voltage-sensitive 7 | 1.71 | 1.50E-02 |
| Gpr157 | G protein-coupled receptor 157 | 1.70 | 2.29E-02 |
| Il18 | interleukin 18 | 1.69 | 2.04E-02 |
| Tent5c | terminal nucleotidyltransferase 5C | 1.69 | 2.94E-02 |
| Il1rn | interleukin 1 receptor antagonist | 1.69 | 5.55E-03 |
| Gzma | granzyme A | 1.68 | 4.40E-02 |
| P2rx4 | purinergic receptor P2X, ligand-gated ion channel 4 | 1.68 | 1.59E-02 |
| Gbp3 | guanylate binding protein 3 | 1.67 | 1.75E-02 |
| Ifih1 | interferon induced with helicase C domain 1 | 1.67 | 8.33E-03 |
| Slfn5 | schlafen 5 | 1.67 | 2.05E-02 |
| Fcgr4 | Fc receptor, IgG, low affinity IV | 1.67 | 4.25E-02 |
| Fcgr1 | Fc receptor, IgG, high affinity I | 1.66 | 6.34E-03 |
| Ldlr | low density lipoprotein receptor | 1.65 | 3.41E-02 |
| Lap3 | leucine aminopeptidase 3 | 1.64 | 3.73E-02 |
| P2ry14 | purinergic receptor P2Y, G-protein coupled, 14 | 1.63 | 4.33E-02 |
| Zfp954 | zinc finger protein 954 | 1.61 | 4.45E-02 |
| Scimp | SLP adaptor and CSK interacting membrane protein | 1.61 | 3.42E-02 |
| Csf1 | colony stimulating factor 1 (macrophage) | 1.60 | 9.21E-03 |
| Slfn3 | schlafen 3 | 1.59 | 4.83E-02 |
| Trim30d | tripartite motif-containing 30D | 1.59 | 8.69E-03 |
| Slc38a6 | solute carrier family 38, member 6 | 1.57 | 3.32E-02 |
| Csf2rb | colony stimulating factor 2 receptor, beta, low-affinity (granulocyte-macrophage) | 1.57 | 4.09E-02 |
| Lgals8 | lectin, galactose binding, soluble 8 | 1.57 | 1.38E-02 |
| Renbp | renin binding protein | 1.55 | 4.40E-02 |
| Cd22 | CD22 antigen | 1.55 | 4.40E-02 |
| Slfn8 | schlafen 8 | 1.55 | 2.30E-02 |
| Il18bp | interleukin 18 binding protein | 1.55 | 2.30E-02 |
| B430306N03Rik | RIKEN cDNA B430306N03 gene | 1.54 | 1.38E-02 |
| Tmem144 | transmembrane protein 144 | 1.54 | 2.87E-02 |
| F13a1 | coagulation factor XIII, A1 subunit | 1.54 | 3.55E-02 |
| Pfkfb3 | 6-phosphofructo-2-kinase/fructose-2,6-biphosphatase 3 | 1.53 | 2.66E-02 |
| Themis2 | thymocyte selection associated family member 2 | 1.53 | 1.81E-02 |
| Dusp3 | dual specificity phosphatase 3 (vaccinia virus phosphatase VH1-related) | 1.51 | 4.14E-02 |
| Osbpl8 | oxysterol binding protein-like 8 | 1.51 | 2.66E-02 |
| Oasl2 | 2'-5' oligoadenylate synthetase-like 2 | 1.51 | 4.07E-02 |
| Chst14 | carbohydrate sulfotransferase 14 | 1.50 | 3.52E-02 |
| Msmo1 | methylsterol monoxygenase 1 | 1.50 | 3.40E-02 |
| Oas1a | 2'-5' oligoadenylate synthetase 1A | 1.49 | 4.81E-02 |
| Nmral1 | NmrA-like family domain containing 1 | 1.48 | 4.16E-02 |
| Fam213b | family with sequence similarity 213, member B | 1.48 | 3.35E-02 |
| Trim30a | tripartite motif-containing 30A | 1.47 | 4.66E-02 |
| Cd40 | CD40 antigen | 1.45 | 4.87E-02 |
| Cipc | CLOCK interacting protein, circadian | 1.45 | 3.42E-02 |
| Samhd1 | SAM domain and HD domain, 1 | 1.45 | 4.02E-02 |

| Gene symbol | Gene name | Absolute fold change (CD122+Macs/cMacs) | Adjusted *p* value |
|---|---|---|---|
| Atp6v1d | ATPase, H+ transporting, lysosomal V1 subunit D | 1.44 | 3.35E-02 |
| Sp100 | nuclear antigen Sp100 | 1.42 | 3.75E-02 |
| St8sia4 | ST8 alpha-N-acetyl-neuraminide alpha-2,8-sialyltransferase 4 | 0.68 | 4.81E-02 |
| Srm | spermidine synthase | 0.67 | 4.81E-02 |
| Nr4a2 | nuclear receptor subfamily 4, group A, member 2 | 0.67 | 3.51E-02 |
| Taf4b | TATA-box binding protein associated factor 4b | 0.67 | 3.51E-02 |
| St3gal6 | ST3 beta-galactoside alpha-2,3-sialyltransferase 6 | 0.67 | 3.42E-02 |
| Serpine2 | serine (or cysteine) peptidase inhibitor, clade E, member 2 | 0.67 | 4.61E-02 |
| St8sia6 | ST8 alpha-N-acetyl-neuraminide alpha-2,8-sialyltransferase 6 | 0.66 | 2.24E-02 |
| Cytip | cytohesin 1 interacting protein | 0.66 | 4.00E-02 |
| Slc25a33 | solute carrier family 25, member 33 | 0.66 | 4.40E-02 |
| AU021092 | expressed sequence AU021092 | 0.66 | 3.11E-02 |
| Plbd1 | phospholipase B domain containing 1 | 0.66 | 3.33E-02 |
| Bend6 | BEN domain containing 6 | 0.66 | 4.99E-02 |
| Mcemp1 | mast cell expressed membrane protein 1 | 0.65 | 3.76E-02 |
| S100a4 | S100 calcium binding protein A4 | 0.64 | 4.29E-02 |
| Rangrf | RAN guanine nucleotide release factor | 0.64 | 4.18E-02 |
| F2r | coagulation factor II (thrombin) receptor | 0.64 | 4.89E-02 |
| Ebpl | emopamil binding protein-like | 0.64 | 1.50E-02 |
| Cers4 | ceramide synthase 4 | 0.63 | 4.18E-02 |
| 1700025G04Rik | RIKEN cDNA 1700025G04 gene | 0.63 | 1.37E-02 |
| Klra2 | killer cell lectin-like receptor, subfamily A, member 2 | 0.63 | 2.58E-02 |
| Spred2 | sprouty-related, EVH1 domain containing 2 | 0.63 | 3.33E-02 |
| Abca9 | ATP-binding cassette, sub-family A (ABC1), member 9 | 0.63 | 4.28E-02 |
| Cd5l | CD5 antigen-like | 0.63 | 4.40E-02 |
| H2-Ob | histocompatibility 2, O region beta locus | 0.63 | 2.22E-02 |
| Atp10d | ATPase, class V, type 10D | 0.63 | 2.12E-02 |
| Ttc28 | tetratricopeptide repeat domain 28 | 0.62 | 4.40E-02 |
| Akr1b8 | aldo-keto reductase family 1, member B8 | 0.62 | 1.93E-02 |
| Hnrnph3 | heterogeneous nuclear ribonucleoprotein H3 | 0.62 | 3.51E-02 |
| Tnfsf13b | tumor necrosis factor (ligand) superfamily, member 13b | 0.61 | 3.94E-02 |
| Cmah | cytidine monophospho-N-acetylneuraminic acid hydroxylase | 0.61 | 3.09E-02 |
| Cd300e | CD300E molecule | 0.61 | 3.42E-02 |
| Ltbp1 | latent transforming growth factor beta binding protein 1 | 0.61 | 1.74E-02 |
| Ifi27 | interferon, alpha-inducible protein 27 | 0.60 | 5.67E-03 |
| Tmem204 | transmembrane protein 204 | 0.60 | 1.75E-02 |
| Hspb1 | heat shock protein 1 | 0.59 | 2.99E-02 |
| Pecam1 | platelet/endothelial cell adhesion molecule 1 | 0.59 | 2.66E-02 |
| Cyr61 | cysteine rich protein 61 | 0.59 | 1.95E-02 |
| Cnn3 | calponin 3, acidic | 0.59 | 1.63E-02 |
| Rbp4 | retinol binding protein 4, plasma | 0.58 | 2.04E-02 |
| Myc | myelocytomatosis oncogene | 0.58 | 3.93E-02 |
| Angpt2 | angiopoietin 2 | 0.58 | 4.81E-02 |
| Padi4 | peptidyl arginine deiminase, type IV | 0.58 | 5.50E-03 |
| Jam2 | junction adhesion molecule 2 | 0.58 | 1.66E-02 |
| Fermt2 | fermitin family member 2 | 0.57 | 3.33E-02 |
| Tcim | transcriptional and immune response regulator | 0.57 | 1.82E-02 |
| Fst | follistatin | 0.57 | 2.35E-02 |
| Fstl1 | follistatin-like 1 | 0.56 | 2.93E-02 |
| Maob | monoamine oxidase B | 0.56 | 2.51E-02 |
| Gm24463 | predicted gene, 24463 | 0.55 | 1.10E-02 |
| Tmem176b | transmembrane protein 176B | 0.55 | 3.44E-02 |
| Olr1 | oxidized low density lipoprotein (lectin-like) receptor 1 | 0.54 | 4.07E-02 |
| Plod2 | procollagen lysine, 2-oxoglutarate 5-dioxygenase 2 | 0.54 | 3.21E-02 |
| H2-Aa | histocompatibility 2, class II antigen A, alpha | 0.53 | 2.28E-03 |
| Ccr2 | chemokine (C-C motif) receptor 2 | 0.53 | 1.14E-02 |
| Gclm | glutamate-cysteine ligase, modifier subunit | 0.53 | 4.69E-03 |
| Tle1 | transducin-like enhancer of split 1 | 0.53 | 2.47E-03 |
| Igfbp3 | insulin-like growth factor binding protein 3 | 0.52 | 1.28E-02 |
| Cxcl14 | chemokine (C-X-C motif) ligand 14 | 0.52 | 3.54E-02 |
| Treml4 | triggering receptor expressed on myeloid cells-like 4 | 0.51 | 2.21E-03 |
| Rgs5 | regulator of G-protein signaling 5 | 0.51 | 2.28E-03 |
| Rdh10 | retinol dehydrogenase 10 (all-trans) | 0.51 | 1.81E-02 |
| Mest | mesoderm specific transcript | 0.51 | 3.42E-02 |

| Gene symbol | Gene name | Absolute fold change (CD122+Macs/cMacs) | Adjusted *p* value |
|---|---|---|---|
| H2-Ab1 | histocompatibility 2, class II antigen A, beta 1 | 0.50 | 2.92E-02 |
| Timd4 | T cell immunoglobulin and mucin domain containing 4 | 0.50 | 5.03E-03 |
| Tmem119 | transmembrane protein 119 | 0.48 | 1.82E-03 |
| Tdo2 | tryptophan 2,3-dioxygenase | 0.48 | 2.41E-03 |
| Clec4b1 | C-type lectin domain family 4, member b1 | 0.47 | 3.87E-02 |
| Igfbp7 | insulin-like growth factor binding protein 7 | 0.47 | 6.34E-03 |
| Tmem176a | transmembrane protein 176A | 0.47 | 2.41E-03 |
| Areg | amphiregulin | 0.47 | 2.55E-02 |
| Cd74 | CD74 antigen (invariant polypeptide of major histocompatibility complex, class II antigen-associated) | 0.46 | 3.67E-03 |
| Plpp1 | phospholipid phosphatase 1 | 0.46 | 1.60E-02 |
| Fpr1 | formyl peptide receptor 1 | 0.46 | 6.34E-03 |
| Sparcl1 | SPARC-like 1 | 0.45 | 6.59E-03 |
| Slpi | secretory leukocyte peptidase inhibitor | 0.45 | 3.94E-02 |
| Mfap5 | microfibrillar associated protein 5 | 0.45 | 2.41E-03 |
| Efemp1 | epidermal growth factor-containing fibulin-like extracellular matrix protein 1 | 0.45 | 1.58E-02 |
| Tgfb2 | transforming growth factor, beta 2 | 0.44 | 6.59E-03 |
| Siglece | sialic acid binding Ig-like lectin E | 0.44 | 7.07E-03 |
| Cd24a | CD24a antigen | 0.42 | 2.05E-02 |
| Hpgd | hydroxyprostaglandin dehydrogenase 15 (NAD) | 0.41 | 6.34E-03 |
| H2-Eb1 | histocompatibility 2, class II antigen E beta | 0.40 | 3.57E-03 |
| F5 | coagulation factor V | 0.40 | 9.12E-03 |
| Plxdc2 | plexin domain containing 2 | 0.40 | 5.52E-04 |
| Retnla | resistin like alpha | 0.38 | 7.82E-03 |
| Vsig4 | V-set and immunoglobulin domain containing 4 | 0.38 | 3.43E-04 |
| Selp | selectin, platelet | 0.34 | 2.41E-03 |
| Fabp7 | fatty acid binding protein 7, brain | 0.30 | 2.76E-03 |
| Fn1 | fibronectin 1 | 0.27 | 5.52E-04 |
| Adgre4 | adhesion G protein-coupled receptor E4 | 0.23 | 2.36E-04 |
| Serpinb10 | serine (or cysteine) peptidase inhibitor, clade B (ovalbumin), member 10 | 0.23 | 1.78E-04 |
| Il6 | interleukin 6 | 0.18 | 3.25E-04 |
| Serpinb2 | serine (or cysteine) peptidase inhibitor, clade B, member 2 | 0.10 | 1.04E-04 |
| Alox15 | arachidonate 15-lipoxygenase | 0.08 | 6.53E-06 |
| Cxcl13 | chemokine (C-X-C motif) ligand 13 | 0.08 | 1.30E-04 |
| Saa3 | serum amyloid A 3 | 0.05 | 4.55E-06 |
| Prg4 | proteoglycan 4 (megakaryocyte stimulating factor, articular superficial zone protein) | 0.04 | 2.46E-05 |

**Supplemental Table 2. Numerous interferon-stimulated genes and cytolytic transcripts distinguish the transcriptome of decidual CD122+Macs from that of decidual cMacs.** Changes in gene expression by microarray between sort-purified CD122+Macs and cMacs. Only those with an adjusted *p* value <0.05 are shown. Absolute fold change of each individual gene refers to level in CD122+Macs relative to level in cMacs. Of note, *Il2rb* (encoding CD122) met threshold unadjusted *p* value but did not meet threshold adjusted *p* value.

#### SUPPLEMENTAL FIGURE 2

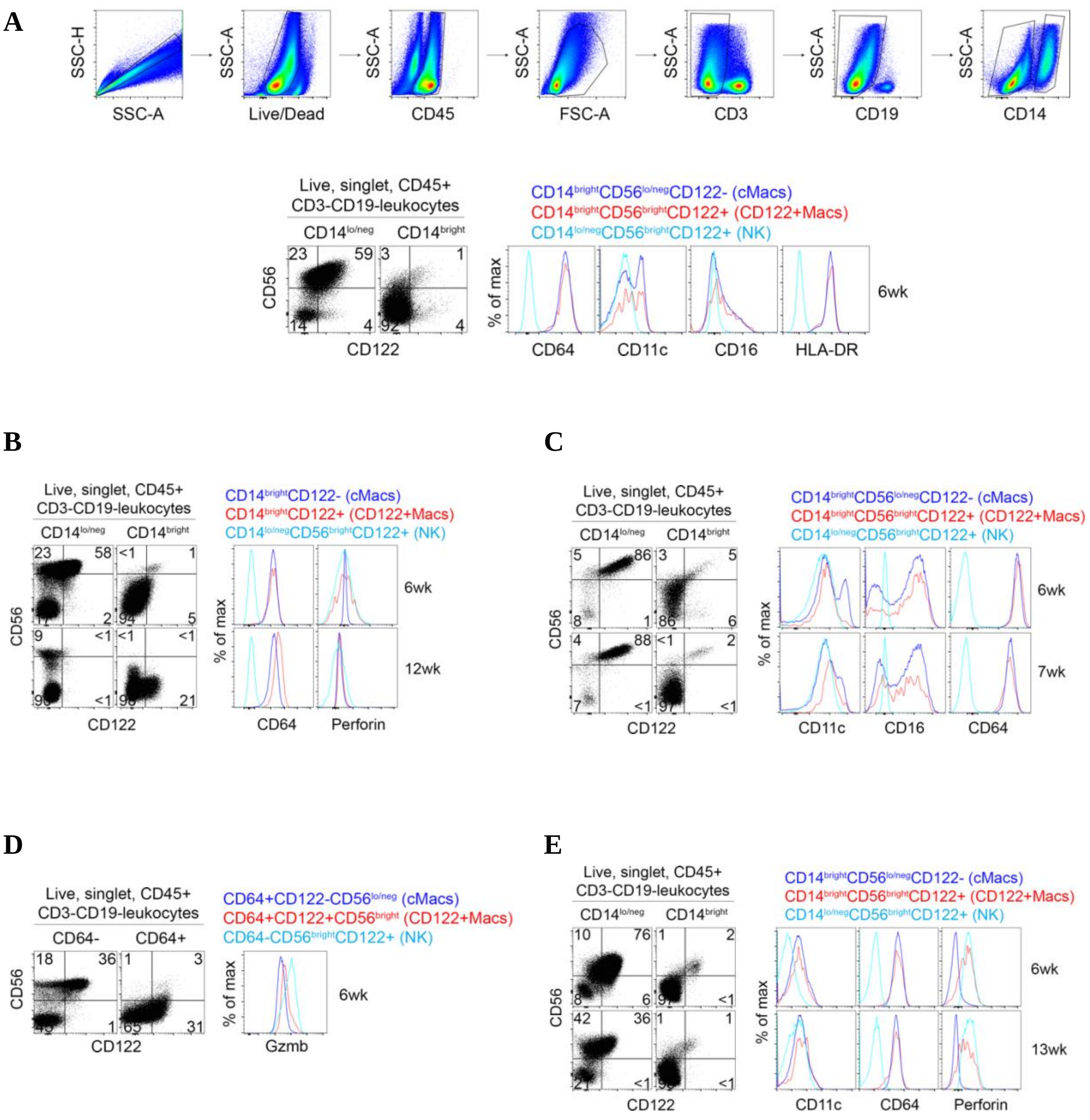


**Supplemental Figure 2. Human CD122+Macs are present in first trimester decidua.** (A-E) Additional examples of human CD122+Macs in first-trimester deciduae analyzed after elective terminations of pregnancy. Shown in (A) is the raw gating scheme to identify human CD122+Macs. Note that human decidual CD122+Macs variably express CD56.

#### SUPPLEMENTAL FIGURE 3

**A**

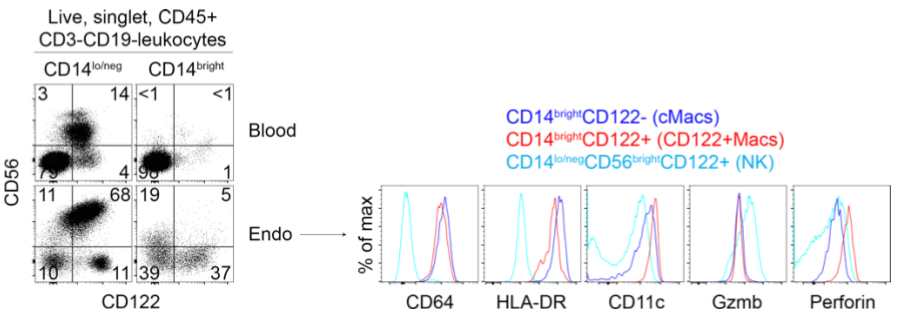


**B**

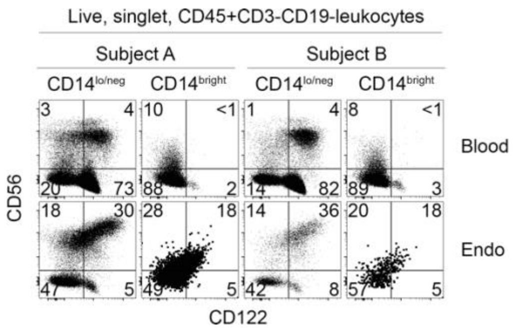


**C**

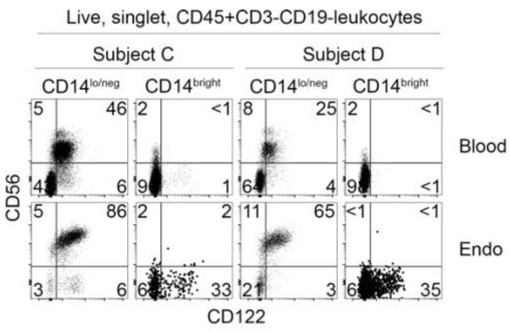


**Supplemental Figure 3. Human CD122+Macs are present in secretory phase endometrium during the implantation window.** (A-C) Additional examples of human CD122+Macs in secretory phase endometrium. Like first-trimester decidual CD122+Macs, endometrial CD122+Macs variably express CD56. Up to two samples were sometimes processed simultaneously. Any samples stained and analyzed on the same day are shown in the same sub-figure, with headings to distinguish each individual subject from one another.

**SUPPLEMENTAL FIGURE 4**

**A**

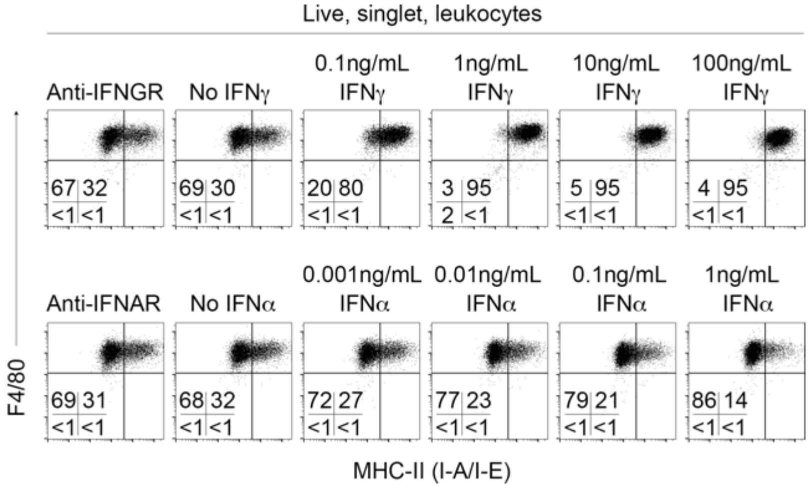


**B**

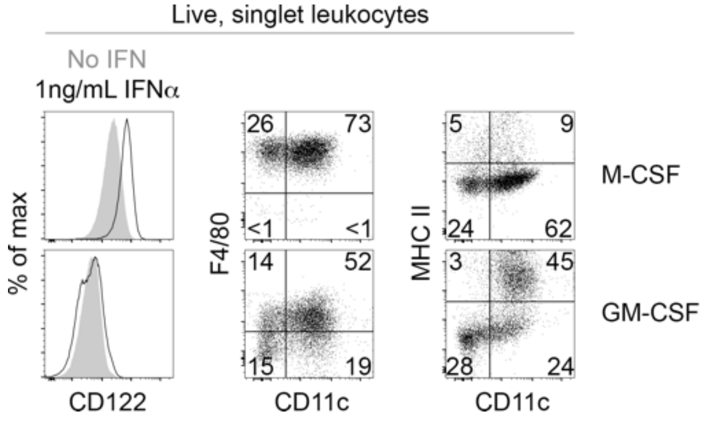


**Supplemental Figure 4. GM-CSF and IFNγ both induce MHCII on BMDMs.** (A) Dose-dependent upregulation of MHC Class II on bone marrow-derived macrophages (BMDMs) cultured with IFNγ (top row). Data are representative of at least 5 independent experiments. (B) BMDM derived with M-CSF are F4/80$_{bright}$ and increase surface CD122 when exposed to exogenous IFNα. BMDM derived with GM-CSF are F4/80$_{int}$, and many express high levels of MHCII. BMDM derived with GM-CSF do not exhibit surface expression of CD122. Data are representative of at least 5 independent experiments.

**SUPPLEMENTAL FIGURE 5**

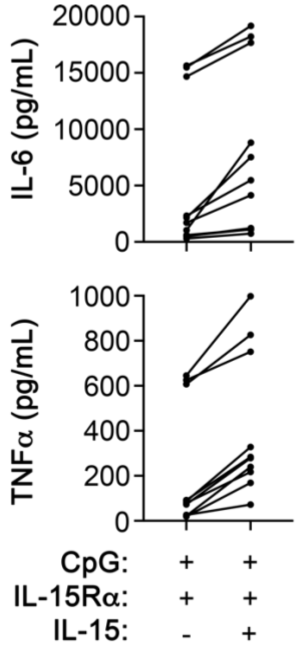


**Supplemental Figure 5. IL-15 costimulation consistently enhances production of IL-6 and TNFα by CD122+BMDMs responding to CpG**. All individual data points, shown as composite bar graphs in Main Figure 7, compiled from 11 biological replicates over 5 independent experiments, with BMDMs in triplicate generated from 1-3 individual mice per experiment.

| | Antigen | Clone | Fluor | Dilution | Manufacturer |
|---|---|---|---|---|---|
| **MU** | CCR2 | 475301 | PE | 1:100 | R&D Systems |
| | CD11b | M1/70 | FITC | 1:200 | BD |
| | CD11c | N418 | PE-Cy7 | 1:200 | Biolegend |
| | CD122 | TM-β1 | BV421 (best), PE | 1:100 | BD |
| | CD19 | 6D5 | APC-Cy7 | 1:400 | Biolegend |
| | CD3 | 17A2 | APC-Cy7 | 1:100 | BIolegend |
| | CD45.1 | A20 | FITC | 1:100 | BD |
| | CD45.2 | 104 | PerCP-Cy5.5 | 1:100 | Biolegend |
| | CD49a | Ha31/8 | PerCP-Cy5.5 | 1:100 | BD |
| | CD49b | DX5 | PE | 1:100 | Biolegend |
| | CD64 | X54-5/7.1 | PE-Cy7 | 1:200 | Biolegend |
| | Eomes | dan11mag | AF488 | 1:200 | Thermo Fisher |
| | F4/80 | BM8 | APC, PE | 1:100 | Biolegend |
| | Ly6C | HK1.4 | AF700, Pacific Blue | 1:400 | BIolegend |
| | Ly6G | 1A8 | APC-Fire 750 | 1:400 | Biolegend |
| | MERTK | 2B10C42 | PE | 1:100 | Biolegend |
| | MHCII | M5/114.15.2 | PerCP-Cy5.5 | 1:200 | BD |
| | NKp46 | 29A1.4 | FITC | 1:100 | Thermo Fisher |
| | T-bet | 4B10 | BV421 | 1:100 | Biolegend |
| **HU** | CD11c | B-ly6 | PE-Cy7 | 1:100 | BD |
| | CD122 | TU27 | PE | 1:100 | Biolegend |
| | CD14 | M5E2 | V450 | 1:100 | BD |
| | CD16 | 3G8 | AF700 | 1:100 | Biolegend |
| | CD19 | HIB19 | BV605 | 1:100 | Biolegend |
| | CD3 | OKT3 | BV785 | 1:100 | Biolegend |
| | CD45 | HI30 | PerCP-Cy5.5 | 1:100 | BD |
| | CD56 | 5.1H11 | BV650 | 1:100 | Biolegend |
| | CD64 | 10.1 | APC Fire 750 | 1:100 | Biolegend |
| | Granzyme B | GB11 | Pacific Blue, FITC | 1:100 | BIolegend |
| | HLA-DR | G46-6 | V500 | 1:100 | BD |
| | Perforin | δG9 | AF647 | 1:100 | BD |

**Supplemental Table 3.** List and description of antibodies used for flow cytometry and cell sorting. MU denotes anti-mouse antibodies, and HU denotes anti-human antibodies.
